# NSD2 links developmental plasticity to mitochondrial function in muscle and lymphocyte differentiation

**DOI:** 10.1101/2025.04.26.648922

**Authors:** Jorge Martínez-Cano, Ana Sagrera, Belén S. Estrada, Sara Cogliati, Nuria Martínez-Martín, César Cobaleda

## Abstract

Developmental processes require a precise regulation of all the aspects of cellular function to successfully achieve the generation of differentiated cells. This regulation does not only encompass gene expression levels activating the developmental programs, but should also attend the demanding energetic needs of the differentiating cells. Epigenetic regulators are essential for establishing cell-type-specific transcriptional programs, but emerging evidence suggests that they also play a more direct role in regulating metabolism. Nsd2 is a Histone-3-Lysine36 mono- and di-methyltransferase key in the development of several cell types, and is involved in human pathologies both by loss- and gain-of-function alterations. Methylation at H3K36 by proteins of the Nsd family has been shown to be crucial in the maintenance of cell identity by being involved in the establishment and sustained expression of cell-type-specific programs. Here, we demonstrate that Nsd2 is essential for coordinating mitochondrial function and metabolic remodeling during B cell activation and muscle progenitor differentiation. In the terminal differentiation of B cells in the germinal center, the absence of Nsd2 results in defective mitochondrial function, characterized by hyperpolarization of the mitochondrial membrane, elevated reactive oxygen species (ROS) production, and reduced oxygen consumption and ATP production. Similarly, Nsd2 deficiency in myoblasts disrupts metabolic reprogramming during muscle differentiation, leading to impaired mitochondrial respiration and structural abnormalities. In both cell types, these alterations correlate with widespread changes in the expression of genes involved in mitochondrial function and cellular metabolism. These findings highlight Nsd2 as a central mediator linking epigenetic regulation with mitochondrial function, underscoring its critical role in coupling transcriptional programs with metabolic adaptation during cell differentiation.

## INTRODUCTION

Cellular differentiation is a highly orchestrated process that requires precise integration of changes in gene expression, metabolism, and epigenetic modifications. The ability of cells to transition from a progenitor stage to a specialized functional cell type is not only dependent on the establishment of transcriptional programs, but also contingent on the right metabolic reprogramming, which ensures that the energetic and biosynthetic demands of differentiation are met. In recent years, accumulating evidence has linked metabolism and mitochondrial function to epigenetics and chromatin regulation, establishing that communication between mitochondria and the nucleus is vital for regulating cellular functions (Matilainen et al., 2017; Zhu et al., 2022). On one side, nuclear-encoded transcription factors, essential for mitochondrial function, are modulated by histone modifications, impacting energy production and oxidative capacity (Scarpulla et al., 2012). Conversely, mitochondrial metabolites are critical for the activity of nuclear epigenetic enzymes, establishing a link between mitochondrial metabolic states and nuclear gene expression (Xu et al., 2011). This bidirectional communication is vital for maintaining cellular homeostasis and it is particularly relevant during differentiation, where metabolic shifts accompany lineage specification from early progenitors (Chandel et al., 2016; Lisowski et al., 2018). Accordingly, dysregulation of these pathways can lead to developmental abnormalities and impaired cell fate transitions, and has significant implications for diseases such as cancer and neuromuscular and immune disorders (Rossmann et al., 2021; Suomalainen and Nunnari, 2024).

Among the epigenetic regulators, the three members of the Nuclear Receptor SET domain-containing family of histone methyltransferases (NSD1, 2 and 3), have been implicated in development, differentiation, and disease (Bennett et al., 2017; Li et al., 2019; Vougiouklakis et al., 2015; Wagner and Carpenter, 2012). They primarily catalyze the di-methylation of histone H3 at lysine 36 (H3K36me2), a modification associated with maintenance of cell identity by contributing to preserve DNA methylation at intergenic regions, antagonizing H3K27me3 and therefore preventing the spreading of repressive chromatin, and by helping to keep enhancers in their existing conformation (Ko et al., 2024; Li et al., 2023; Li et al., 2022; Pashos et al., 2025; Sengupta et al., 2021; Shao et al., 2024; Zhao et al., 2021). The three *NSD* paralogs are each catalytically dominant in different tissue types and, therefore, their individual loss is only noticeable in the cell types or developmental stages depending exclusively or mainly on the respective gene affected. NSD2 (also known as WHSC1 or MMSET) is a representative example of the role of epigenetic regulators in both normal and pathological development, since alterations in its function, either by loss- or gain-of-function, have severe consequences for the organism. Loss-of-function mutations in *NSD2* are involved in Wolf-Hirschhorn Syndrome (WHS, hence the other name of *NSD2*, “Wolf-Hirschhorn Syndrome Candidate 1”, *WHSC1*), a rare disease (1 per 20,000 births) caused by the loss of genetic material in the “p” arm of chromosome 4 (Battaglia et al., 2015; Campos-Sanchez et al., 2019; Nevado et al., 2020), where the affected patients suffer from a plethora of conditions affecting many organs. The major causes of morbidity and mortality among WHS patients are seizures, low muscular tone and immunodeficiency (Zollino and Doronzio, 2018). The latter includes an impaired response to lipopolysaccharides, and IgA and IgG2 deficiencies, and it causes an increased susceptibility to infections (Hanley-Lopez et al., 1998). Research from our group and others has identified the hemizygous loss of *NSD2* as the main responsible for immunodeficiency in WHS (Campos-Sanchez et al., 2017; Campos-Sanchez et al., 2019; Dobenecker et al., 2020; Long et al., 2020; Nguyen et al., 2017; Nimura et al., 2009; Pei et al., 2013). Indeed, *Nsd2* was shown to be essential for the normal differentiation and function of B lymphocytes at several developmental stages; first, both specification and commitment to the B cell lineage were impaired in *Nsd2^-/-^* lymphoid progenitors, which exhibited significantly reduced expression of transcription factors Ebf1 and Pax5, essential for the earliest stages of B cell development. At later stages, a second functional and developmental impairment of *Nsd2^-/-^* B cells took place during germinal center (GC) formation, where class switch recombination (CSR) was significantly reduced in an *Nsd2*-dose-dependent manner (Campos-Sanchez et al., 2017). This impairment was accompanied by aberrant proliferation of *Nsd2*-deficient B cells, correlating with an elevated replicative stress associated with accumulation of DNA damage. Another aspect that supports the functional relevance of NSD2 in lymphocyte development is its participation in B-cell-associated human malignancies, like multiple myeloma (MM, hence the third name of *NSD2*, “Multiple Myeloma SET domain”, *MMSET*), where it exhibits oncogenic activity when overexpressed in plasma cells harboring the t(4;14)(p16.3;q32.3) translocation (Larrayoz et al., 2023; Stec et al., 1998), and childhood B-cell acute lymphoblastic leukemias, where activating gain-of-function mutations in *NSD2* collaborate in the progression of the disease (Jaffe et al., 2013; Studd et al., 2021). Besides hematopoietic tumors, NSD2 has also been involved by several mechanisms in the pathogenesis of different solid malignancies (Aytes et al., 2018; Ezponda et al., 2013; Parolia et al., 2024; Sengupta et al., 2021), highlighting its important role in the specification of cell fate.

Besides immunodeficiency, neuromuscular problems (mainly epileptic convulsions and low muscular tone) are the other main complication that WHS patients face, and cause them significant morbidity, developmental delay and disabilities (Correa et al., 2022; Correa et al., 2018). It has been suggested that these neuromuscular defects are the product of a synergistic relationship among several candidate genes specifically related to this type of disorder (Correa et al., 2022; Correa et al., 2018; Ho et al., 2018; Zollino and Doronzio, 2018), and they have been associated with mitochondrial malfunction and metabolic alterations (Hart et al., 2014). Genotype-phenotype correlation studies in WHS patients have identified a genomic region in 4p susceptible to accumulating mutations that may explain the onset of seizures (Seizures Susceptibility Region, SSR) (Correa et al., 2018; Zollino and Doronzio, 2018). In this region, immediately adjacent to *NSD2* in 4p16.3, is located *LETM1*, encoding for a calcium transporter located in the inner mitochondrial membrane; because of this, and also due to the phenotype of *Letm1^-/-^* mice, haploinsufficiency of this mitochondrial protein has been considered the main responsible for the neuromuscular pathologies affecting WHS patients (Hart et al., 2014). However, in the case of WHS, the association of *LETM1* haploinsufficiency with seizures remains contentious due to the existence of clinical cases of patients with *LETM1* deletions but lacking seizures and vice versa (Correa et al., 2022; Maas et al., 2008; Zollino et al., 2014). Nowadays it is increasingly clear that mitochondrial dysfunction is a significant cause of seizures in various pathologies presenting them, potentially resulting from impaired mitochondrial function, abnormal ion fluxes, or elevated mitochondrial reactive oxygen species (mROS) production (Folbergrova and Kunz, 2012). Interestingly, despite being part of the Seizures Susceptibility Region, *NSD2*’s consideration as a candidate seizure gene remains debated today due to insufficient evidence linking its function to altered mitochondrial function, unlike other SSR genes such as *LETM1* (Correa et al., 2018; Zollino and Doronzio, 2018).

Therefore, given all these evidences, and considering that neuromuscular complications, seizures, hypotonia, and failure to thrive, along with immunodeficiencies, are the key characteristics of Wolf-Hirschhorn Syndrome, we aimed to further investigate whether the loss of an epigenetic modifier such as NSD2, involved in the fine control of cell fate changes, could contribute to these phenotypes by disrupting mitochondrial function.

## RESULTS

### Deregulated expression of mitochondrial and metabolic genes in activated *Nsd2^-/-^* B cells

We have previously described how terminal differentiation of B cells and formation of the germinal center reaction are impaired upon reduction of the levels of *Nsd2*, in a dose-dependent manner (Campos-Sanchez et al., 2017). RNA-seq analyses of differential gene expression comparing WT and *Nsd2^-/-^* activated B cells has shown that the effects of the loss of Nsd2 are pleiotropic and affect a plethora of cellular functions, including interference with cell cycle and an impairment in DNA replication efficiency (Campos-Sanchez et al., 2017). Recent research has shown that regulation of metabolic pathways in activated immune cells plays a critical role in fate decision and function during B cell differentiation (Ganeshan and Chawla, 2014; Jang et al., 2015; Jung et al., 2019; Pearce and Pearce, 2013; Waters et al., 2018; Weisel et al., 2020) and, while naïve B cells are quiescent, GC B cells and activated B cells are highly proliferative and metabolically active (Haniuda et al., 2020; Iborra-Pernichi et al., 2024; Mendoza et al., 2018; Weisel et al., 2020). Therefore, we carried out a reanalysis of the RNA-seq data with this focus in mind. This reanalysis revealed that *Nsd2* deficiency led to a deregulation of the expression of a plethora of mitochondrial and metabolic genes in *ex vivo* activated *Nsd2^-/-^* B cells (Figure 1). Pathway analysis using both GO and KEGG databases showed significant alterations in key metabolic processes involved in cellular development (Figure 1A) as exemplified by the big differences in the expression profiles of the KEGG “Metabolic Pathways” gene set (Figure 1B). Regarding more functionally restricted pathways, strong differences in gene expression were found for both cytoplasmic and mitochondrial nuclearly encoded ribosomal protein genes, mitochondrial inner membrane respiratory complex genes and genes involved in general in oxidative phosphorylation (Figure 1C-F). All these data indicate that Nsd2 is necessary for the correct control of the metabolic rewiring and mitochondrial remodeling required for successful B cell activation and class switching.

**Figure 1.**
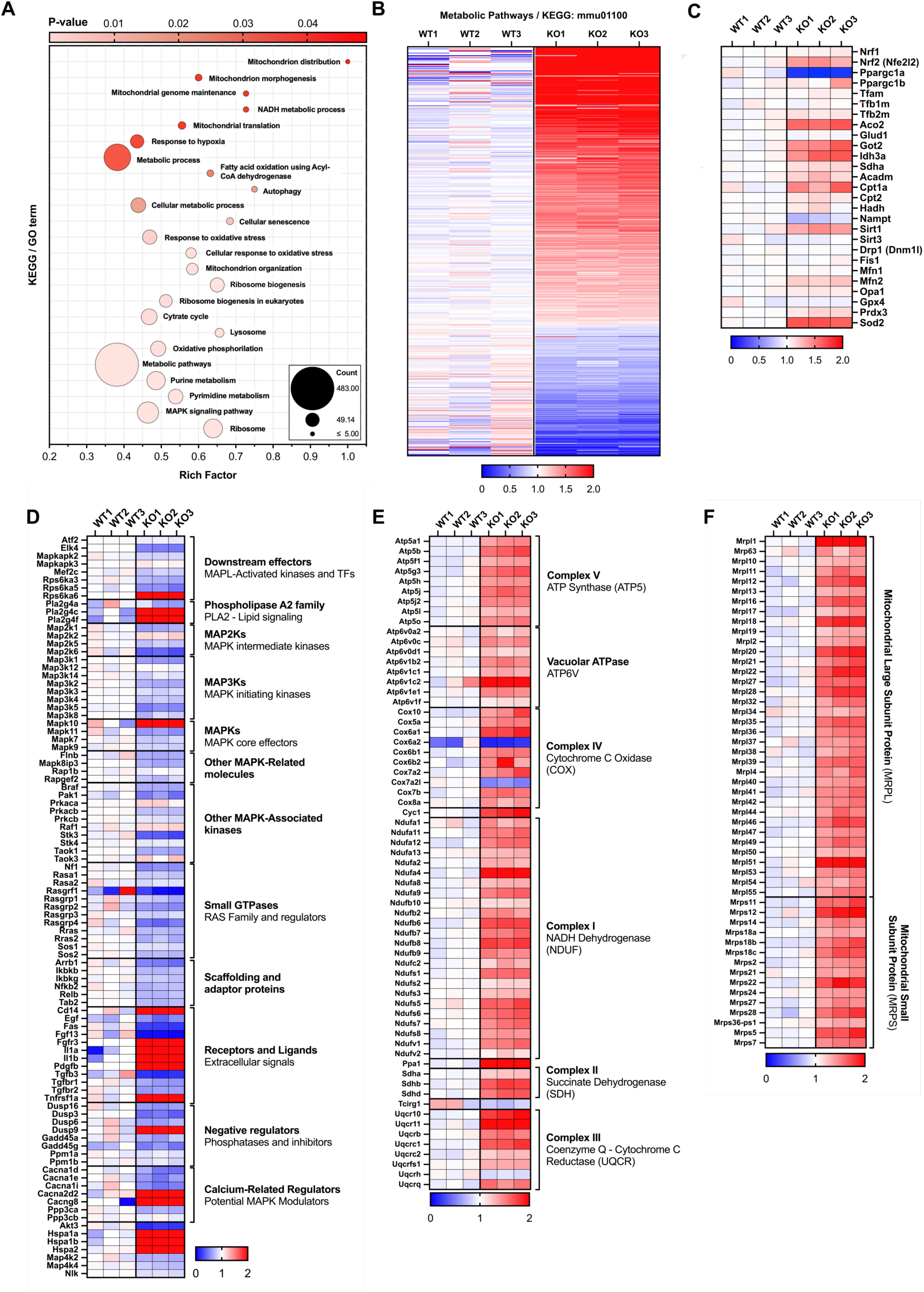
Deregulated expression of mitochondrial and metabolic genes in activated *Nsd2^-/-^*B lymphocytes. Genes differentially expressed when comparing RNA-seq data obtained from *ex vivo* activated splenic B cells of WT or *Nsd2^-/-^* genotypes. **A)** Bubble plot showing the functional enrichment analysis of differentially expressed genes (DEGs) in activated B lymphocytes (WT vs. *Nsd2^-/-^*). The main altered metabolic pathways are shown, selected from either Gene Ontology (GO) or Kyoto Encyclopedia of Genes and Genomes (KEGG) databases. Rich Factor=Total number of DEGs in a given pathway/Total number of genes in that pathway. Circle diameter= Total number of Genes in a pathway. *p*-values according to the color scale on the top. **B)** Heatmap of gene expression profiles for the “*Metabolic Pathways*” gene set from the KEGG database (KEGG:mmu01100). Three independent biological replicates are shown for WT (left three columns) and *Nsd2^-/-^* (right three columns) cells. The normalized Reads Per Kilobase Million (RPKM) values, relative to WT replicates, are displayed for the 483 genes in the set. The color scale is indicated on the bottom horizontal axis. **C)** Heatmap of gene expression profiles of selected key genes involved in mitochondrial genesis and function. **D)** Heatmap of gene expression profiles of key deregulated genes selected from the KEGG:mmu01100 geneset involved in the indicated cellular functions. **E)** Heatmap of gene expression profiles for selected nuclearly encoded genes codifying proteins of the indicated Complexes of the respiratory chain of the inner mitochondrial membrane. **F)** Heatmap of gene expression profiles for selected nuclearly encoded genes codifying protein components of either cytosolic or mitochondrial ribosomes. Three independent biological replicates are shown for WT (left three columns) and *Nsd2^-/-^* (right three columns) cells. The normalized Reads Per Kilobase Million (RPKM) values, relative to WT replicates, are displayed. The color scale is indicated on the bottom horizontal axis.

### Defective Nsd2 function causes mitochondrial membrane hyperpolarization and ROS accumulation in *ex vivo* and *in vivo* stimulated B cells

Next, we wanted to determine how these changes in gene regulation impacted actual mitochondrial function. The maintenance of the correct mitochondrial membrane potential (ΔΨm) is essential for mitochondrial function, and its value is one of the key indicators of such function. Therefore, as a first way of assessing the existence of mitochondrial malfunctions in the absence of Nsd2, we monitored this parameter in WT *Nsd2^+/-^*, and *Nsd2^-/-^* splenic B cells that had been stimulated with LPS+IL4 for 72 h as previously described (Campos-Sanchez et al., 2017), recapitulating the known defects in class switch recombination (CSR) associated to the progressive decrease of dosage of *Nsd2* (Figure 2A-B). We labelled the cells with two different mitochondria-targeted fluorescent probes, MitoTracker Deep Red (MTDR) and TMRM, whose fluorescence intensity is directly proportional to ΔΨm. Mean fluorescence intensity (MFI) histograms obtained after FACS analysis showed (Figure 2C-D) that *Nsd2^+/-^* and, significantly, *Nsd2^-/-^* cells present an increase in the signal for both probes, both in the total culture, and also when switched (IgG1^+^) and unswitched (IgG1^-^) B cells are separately analyzed, although the effect on the cells that managed to switch is less noticeable. To confirm that this effect was not only an artifact appearing in *ex vivo* stimulated cells, *Nsd2^-/-^* and WT mice were stimulated by intraperitoneal injection of Sheep Red Blood Cells (SRBCs), to promote the formation of germinal centers (GCs) *in vivo* through a T-cell-mediated antigenic stimulation (Figure 3A-B), and the initiation of the process was characterized by performing flow cytometric analysis of the spleens 5 days after SRBC injection; once more, it was confirmed that the efficiency of the process is significantly decreased in *Nsd2^-/-^* animals (Campos-Sanchez et al., 2017) (Figure 3A-B). Using MTDR (Figure 3C-D) and TMRM (Figure 3E-F) to characterize mitochondrial status, both probes showed a significant signal increase in *Nsd2^-/-^* cells, indicating that the lack of Nsd2 function leads to an increase in mitochondrial membrane potential, supporting the need of this gene for correct mitochondrial function during B cell activation. On the other hand, the presence of cells with low or null values of MitoTracker Deep Red is an indication of mitochondrial dysfunction, typically associated with loss of mitochondrial membrane potential, early apoptosis, or cellular stress; Supplementary Figure 1 shows the presence of increased percentages of MitoTracker Deep Red^LOW^ cells in *Nsd2^-/-^* activated B cells, both *ex vivo* and *in vivo*.

**Figure 2.**
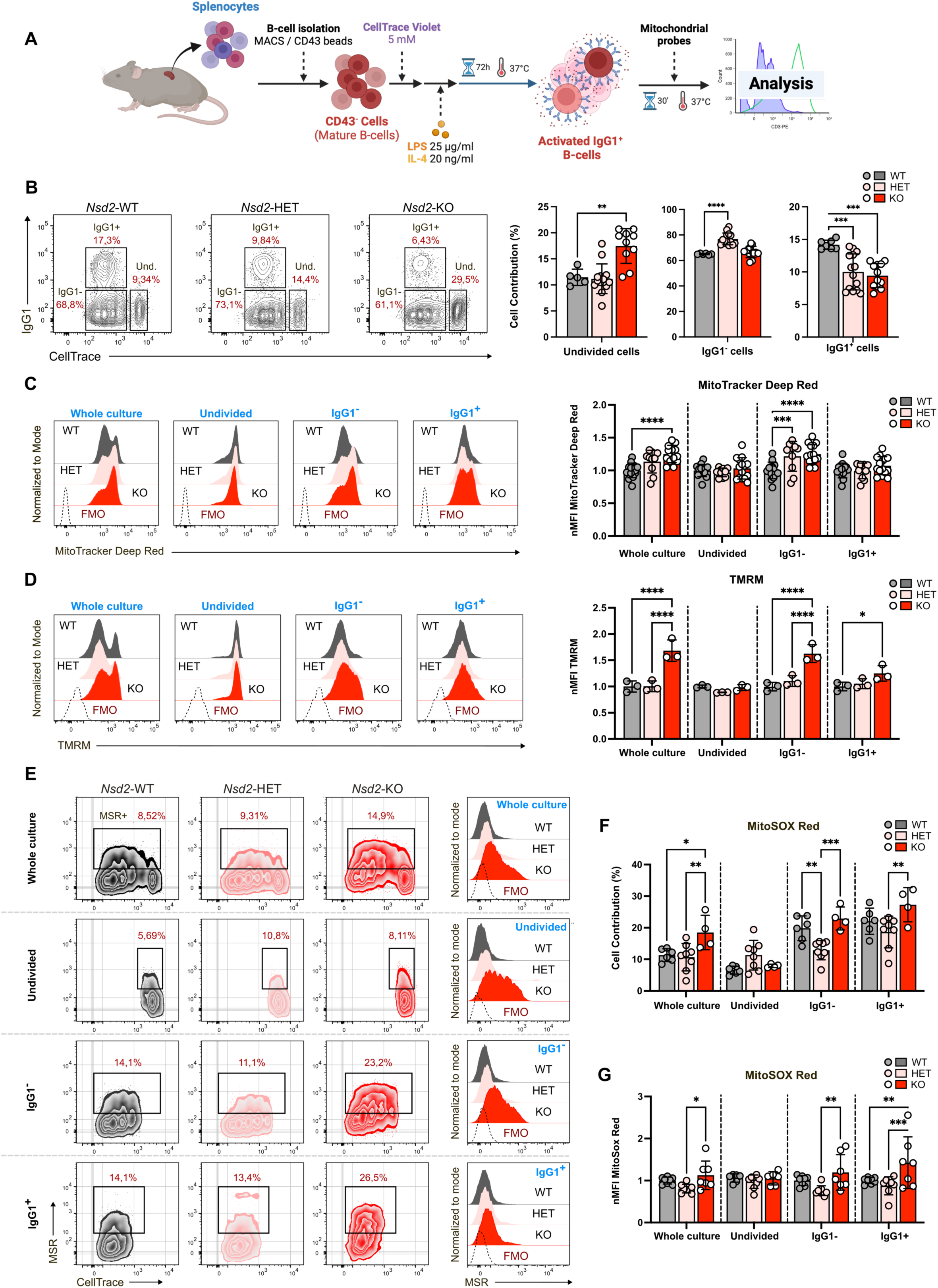
Mitochondrial membrane hyperpolarization and ROS accumulation in *ex vivo* activated *Nsd2^+/-^* or *Nsd2^-/-^*B lymphocytes from spleen. **A)** Schematics of the *ex vivo* B cell activation experimental procedure. **B)** Representative FACS analysis to determine proliferation (as measured by CellTrace dilution) and class switching (as determined by appearance of IgG1 expression) of *ex vivo* stimulated B cells from the spleen, showing the impairment in both parameters –accumulation of undivided cells and decreased percentage of IgG1^+^ cells– with the decrease of *Nsd2* dosage. Bar graphs on the right summarize the results obtained. Biological replicates were analyzed from 3 independent experiments: WT (n=3), HET (n=3), and KO (n=5), with 2–3 technical replicates per measurement. **C)** Histogram overlay representation showing the frequency distribution of MitoTracker Deep Red fluorescence intensity in four population subsets: whole culture, undivided cells, IgG1⁺ cells, and IgG1⁻ cells. Solid dark grey, WT; solid pink, *Nsd2^+/-^* (HET); solid red, *Nsd2^-/-^* (KO); Hollow dashed line, *Fluorescence Minus One* cytometry control (FMO). Bar graphs on the right represent the Mean Fluorescence Intensity of MitoTracker Deep Red, normalized to the wild-type (WT) control from the corresponding experiment. **D)** Histogram overlay representation showing the frequency distribution of the fluorescence intensity of TMRM in the aforementioned four population subsets. Bar graphs on the right represent the Mean Fluorescence Intensity of TMRM, normalized to the wild-type (WT) control from the corresponding experiment. **E)** Representative FACS plots showing MitoSOX Red (MSR) staining across the aforementioned four cellular subsets. Overlaid histograms compare fluorescence emission between unstained FMO controls (dashed black line) and MitoSOX Red–treated samples of the indicated genotypes, with the same color code indicated above highlighting differences in mitochondrial superoxide levels. **F)** Bar graph showing the proportion of MitoSOX Red–positive (MSR⁺) cells in each compartment (total culture, undivided cells, IgG1⁺ cells, and IgG1⁻ cells) for each genotype. **G)** Bar graphs representing the Mean Fluorescence Intensity of MitoSox Red-positive cells, normalized to the wild-type (WT) control from the corresponding experiment. Biological replicates were analyzed from 3 independent experiments: WT (n=3), HET (n=3), and KO (n=5), with 2–3 technical replicates per measurement. Statistical significance was assessed using an unpaired Student’s t test. *(p<0.05), **(p<0.005), ***(p<0.0005), ****(p<0.0001).

**Figure 3.**
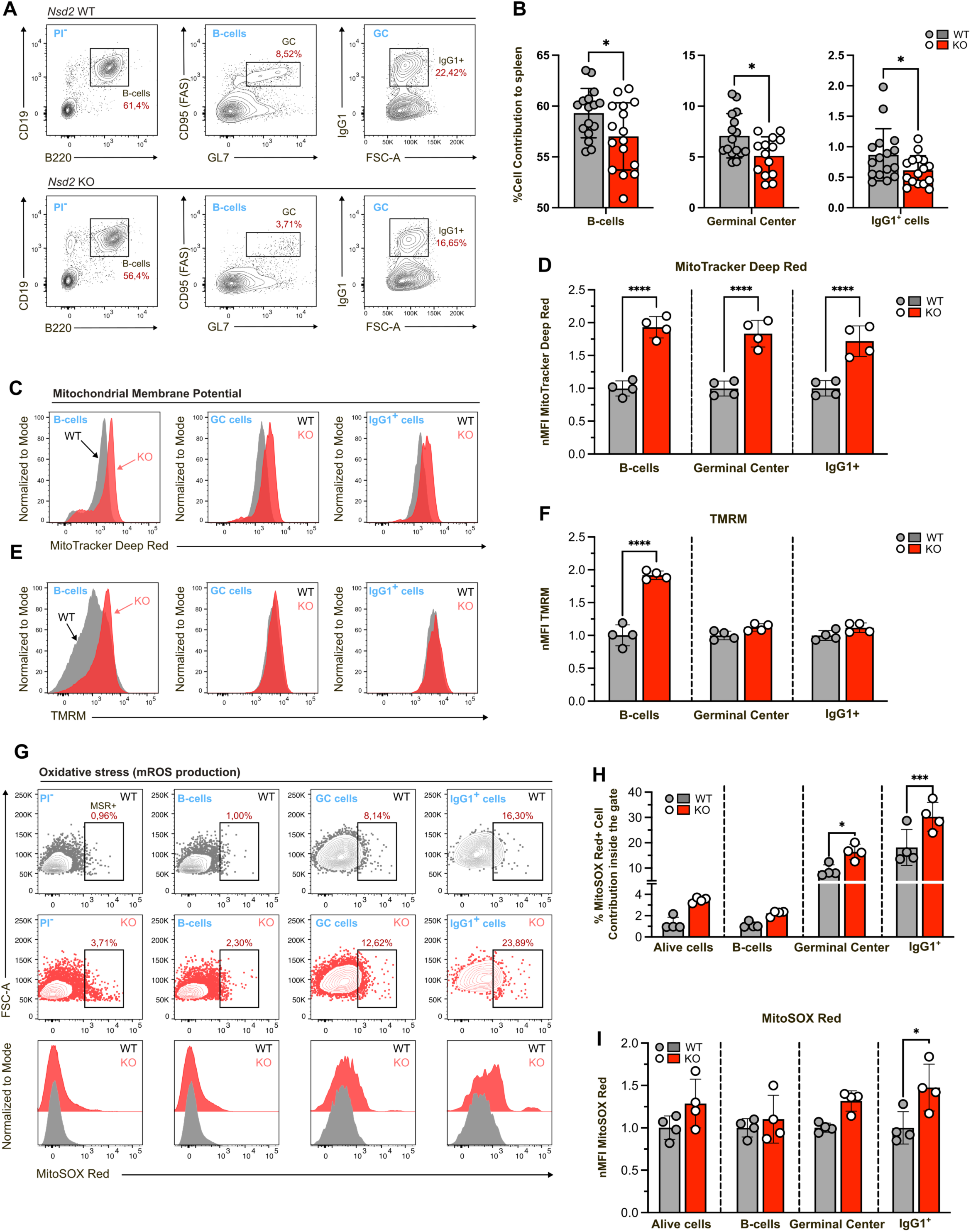
Mitochondrial membrane hyperpolarization and ROS accumulation in *in vivo* Germinal Center *Nsd2^-/-^* B lymphocytes. Mice of the indicated genotypes where intraperitoneally injected with Sheep Red Blood Cells (SRBC) to induce a T cell-dependent immune reaction, and Germinal Center (GC) formation was analyzed in the spleens 5 days after injection. **A)** Representative FACS plots of GC reaction in injected mice of the indicated genotypes (2 WT and 2 *Nsd2^-/-^* –KO– animals, 2 experimental replicates each). Blue labels indicate the backgating of each plot. **B)** Bar graphs showing the population frequencies, as determined by FACS analysis as stated in the previous panel, of the subsets indicated below each plot. Data represent mean ± SD from 2 WT and 2 *Nsd2^-/-^* animals, with 2–4 technical replicates per mouse. **C)** Histogram overlay showing the frequency distribution of MitoTracker Deep Red fluorescence intensity of the cells in the indicated gates (blue labels) in the spleen of SRBC-injected mice of the specified genotypes [solid dark grey, WT; solid red, *Nsd2^-/-^* (KO)]. **D)** Bar graphs representing the Mean Fluorescence Intensity of MitoTracker Deep Red, normalized to the wild-type (WT) control. **E)** Histogram overlay showing the frequency distribution of TMRM fluorescence intensity of the cells in the indicated gates (blue labels) in the spleen of SRBC-injected mice of the specified genotypes [solid dark grey, WT; solid red, *Nsd2^-/-^* (KO)]. **F)** Bar graphs representing the Mean Fluorescence Intensity of TMRM, normalized to the wild-type (WT) control. **G)** Representative FACS plots showing MitoSOX Red (MSR) staining across the four gated subpopulations indicated in blue in the spleen of SRBC-injected mice of the specified genotypes [solid dark grey, WT; solid red, *Nsd2^-/-^* (KO)]. Histogram overlay (lower row) shows the frequency distribution of MitoSOX Red fluorescence intensity of the cells in the same gates as the plots above. **H)** Bar graph showing the proportion of MitoSOX Red–positive (MSR⁺) cells in each gated subpopulation for each genotype. **I)** Bar graph showing the Mean Fluorescence Intensity of MitoSox Red-positive cells, normalized to the wild-type (WT) control. Data represent mean ± SD from 2 WT and 2 *Nsd2^-/-^* animals, with 2–4 technical replicates per mouse. The Kolmogorov-Smirnov test was used to verify the normal distribution of the data. Statistical significance was determined using the independent Student’s t-test. *(p<0.05), **(p<0.005), ***(p<0.0005), ****(p<0.0001).

Alterations in mitochondrial function are frequently accompanied by the appearance of oxidative stress (Elfawy and Das, 2019; Pagano et al., 2014), since a significant part of the levels of reactive oxygen species (ROS) are generated as byproducts of respiration, and one of the main sources of oxidative stress is the overproduction of mitochondrial ROS, being the superoxide anion (O2•−) the most abundant one. Therefore, as a measure of mitochondrial stress, we determined the cellular concentration of superoxide by staining *ex vivo* activated B cells with the fluorescent probe MitoSOX Red (MSR). Figures 2E-G show the contribution to the culture of the stimulated cells that produce excessive levels of mitochondrial superoxide anion (MSR^+^), for both the total culture, or when analyzing undivided, IgG1^+^, or IgG1^-^ cells separately. In general, there is a significant increase in MSR^+^ cells in both *Nsd2^+/-^* and *Nsd2^-/-^* cultures in comparison with those from WT cells, both for the total culture and when distinguishing between switched and unswitched cells (Figure 2F) The analyses of total MitoSOX Red fluorescence levels (Figure 2G) show significantly higher mean fluorescence intensity (MFI) in the IgG1^+^ compartment of *Nsd2^-/-^* cells, suggesting that not only is there a larger accumulation of MSR+ cells within the IgG1^+^ compartment but, also, that they generate larger amounts of superoxide anion, therefore indicating that in the absence of Nsd2 there is an increase in the levels of intracellular ROS. To extend these *ex vivo* results to a more physiological setting, *Nsd2^-/-^* and WT mice were intraperitoneally injected with SRBCs, and splenic GC B cells were analyzed 5 days after injection (Figure 3G-I) In these conditions, again *Nsd2^-/-^* B cells presented a significant increase in the MitoSOX Red signal in both IgG1^+^ and IgG1^-^ GC populations (Figure 3G-I).

In summary, both the *ex vivo* and *in vivo* data from splenic activated B cells confirm that decreased levels of Nsd2 cause mitochondrial membrane hyperpolarization and ROS accumulation, suggesting the existence of Nsd2-dependent alterations in mitochondrial membrane potential.

### Mitochondrial mass, number and morphology are altered in *Nsd2^-/-^* activated B cells

Mitochondrial function and protein composition are closely related to the organelle’s ultrastructure, in such a way that functional mitochondrial defects often correlate with changes in its morphology (Campello and Scorrano, 2010; Miyazono et al., 2018). The MitoTracker Green (MTG) probe specifically accumulates in the mitochondrial matrix, so the fluorescence intensity of MTG-stained cells is directly proportional to their total mitochondrial mass. Figures 4A-B show a representative FACS plot of MTG fluorescence emission in *ex vivo* stimulated mature B cells of the three genotypes (WT, *Nsd2^+/-^* and *Nsd2^-/-^*). Although there was a noticeable increase in the MTG fluorescence levels in *Nsd2^+/-^* and *Nsd2^-/-^* cells in comparison to the control, it was only statistically significant in the unswitched IgG1^-^ compartment. Taking advantage of the dilution of CFSE fluorescent labelling with each cell division, the population could be subdivided into its successive generations in culture (Supplementary Figure 2A-B); this allows detecting an upwards trend in MTG levels in the IgG1^+^ subpopulation of *Nsd2^+/-^* and *Nsd2^-/-^* cells as they divide, in comparison to WT ones (Supplementary Figure 2C-H). These results were confirmed in splenocytes 5 days after SRBC immunization, where we could see that, also *in vivo*, the changes in MitoTracker Green fluorescence in *Nsd2^-/-^* total B cells, in comparison to WT cells, showed a tendency towards an increase in mitochondrial mass (Figure 4C-D).

**Figure 4.**
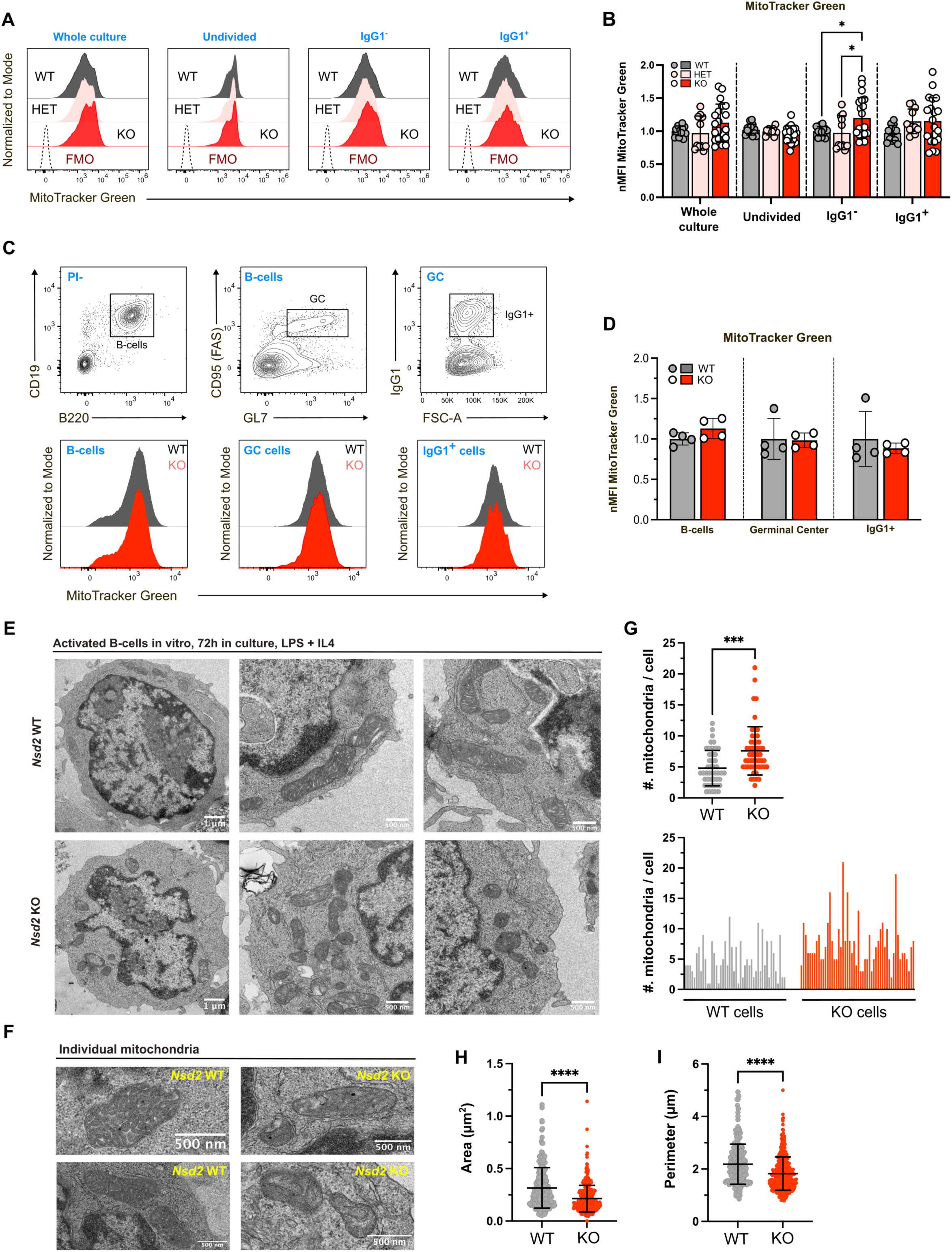
Alterations in mitochondrial mass, number and morphology in activated *Nsd2^-/-^* B cells. **A)** Histogram overlay representation showing the frequency distribution of MitoTracker Green fluorescence intensity in four population subsets (whole culture, undivided cells, IgG1^-^ cells, and IgG1^+^ cells) of *ex vivo* activated B cells from the spleens of mice of the indicated genotypes [Solid dark grey, WT; solid pink, *Nsd2^+/-^* (HET); solid red, *Nsd2^-/-^* (KO); Hollow dashed line, *Fluorescence Minus One* cytometry control (FMO)]. **B)** Bar graph representing the Mean Fluorescence Intensity of MitoTracker Green, normalized to the wild-type (WT) control from the corresponding experiment,as shown in panel A. Biological replicates were analyzed from 3 independent experiments: WT (n=3), HET (n=3), and KO (n=5), with 2–3 technical replicates per measurement. Statistical significance was assessed using an unpaired Student’s t test. **C)** Histogram overlay (lower row) showing the frequency distribution of MitoTracker Green fluorescence intensity of the cells in the indicated gates (blue labels in upper row) in the spleen of SRBC-injected mice of the specified genotypes, 5 days after injection [solid dark grey, WT; solid red, *Nsd2^-/-^* (KO)]. **D)** Bar graphs representing the Mean Fluorescence Intensity of MitoTracker Green, normalized to the wild-type (WT) control for the experiments shown in panel C. Biological replicates were analyzed from 2 independent experiments: WT (n=2) and KO (n=2), with 2 technical replicates per measurement. Statistical significance was assessed using an unpaired Student’s t test. **E,F)** Representative transmission electron microscopy images depicting the mitochondrial structure of activated WT or *Nsd2^-/-^* B lymphocytes at different magnifications. **G-I)** Scatter plots representing the quantification of TEM micrographs parameters in WT vs. *Nsd2* KO cells; G) number of mitochondria per cell, H) cell area (in µm²), and I) cell perimeter (in µm). The Kolmogorov-Smirnov test was used to verify the normal distribution of the data. Statistical significance was determined using the independent Student’s t-test. *(p<0.05), **(p<0.005), ***(p<0.0005).

To better characterize mitochondrial morphology in the absence of *Nsd2*, we decided to ascertain whether mitochondrial ultrastructure was affected by using transmission electron microscopy (TEM) to characterize *ex vivo* stimulated *Nsd2^-/-^* B cells. Figures 4E-F show representative micrographs of activated *Nsd2^-/-^* B cells, where it can be appreciated that, in comparison with WT cells, their mitochondria present some subtle alterations, especially in the definition and density of the cristae inside the organelle; furthermore, *Nsd2^-/-^* mitochondria were smaller and were present in a larger number. A detailed quantification of these parameters (Figure 4G-I) confirmed that *Nsd2^-/-^* lymphocytes present more mitochondria (an average of 7.60 mitochondria/cell) than WT ones (4.80 mitochondria/lymphocyte), and that mitochondria are smaller (0.214 μm^2^) in *Nsd2^-/-^* cells than in WT ones (0.316 μm^2^). This was confirmed by the values of the perimeter of the mitochondria in *Nsd2^-/-^* lymphocytes (1.823 μm) in comparison to WT ones (2.184 μm). Therefore, the absence of Nsd2 correlates to an increase in the number of mitochondria per cell, but these mitochondria are smaller in size; these results correlate with the aforementioned increase in total mitochondrial mass that could be detected using the MitoTracker Green probe (Figure 4A-D).

### Mitochondrial respiration is compromised in *ex vivo* stimulated *Nsd2^-/-^* B cells

Since the results obtained using TMRM and MitoTracker DeepRed probes indicated alterations in mitochondrial membrane potential in the absence of *Nsd2* and, given that conservation of membrane potential is essential for ATP synthesis, we decided to study the cellular respiration of *ex vivo*-activated *Nsd2^-/-^* B cells using extracellular flux analysis (SeaHorse^TM^). We could observe that, in the absence of Nsd2, 72 hours after *ex vivo* stimulation, B cells have considerably lower oxygen consumption rates (OCR) than those recorded for WT cultures, in all the conditions measured (Figure 5A-B and Supplementary Figure 3). All the respiratory parameters were severely affected in the absence of Nsd2, indicating the cell’s reduced ability to cope with conditions of energetic stress and high demand for energy. This weakness of *Nsd2^-/-^* cells upon activation is exacerbated in a time-dependent manner (Figure 5C-D and Supplementary Figure 3), in such a way that, by 90h in culture, KO lymphocytes present with an absence of oxygen consumption for ATP production and a null spare respiratory capacity. These findings further confirm that Nsd2 deficiency impairs mitochondrial respiratory efficiency, limiting the cell’s capacity to meet increased energetic requirements or respond effectively to mitochondrial stress. A similar conclusion can be drawn from the assessment of coupling efficiency, a parameter that indicates the proportion of oxygen consumption dedicated to ATP synthesis compared to the amount driving proton leak (i.e., the fraction of basal mitochondrial OCR utilized for ATP production) (Figure 5E, and data from the ATP-linked OCR at 90h in the bar graph in Figure 4-D); at 72h, although there is a reduction in the efficiency of *Nsd2^-/-^* cells when compared with that from WT cells, the difference is not very pronounced. However, at 90h (Figure 5E), the coupling efficiency measured in the *Nsd2^-/-^* culture is zero while, in the case of WT cells, only a small decrease is observed respect to the value at 72h.

**Figure 5.**
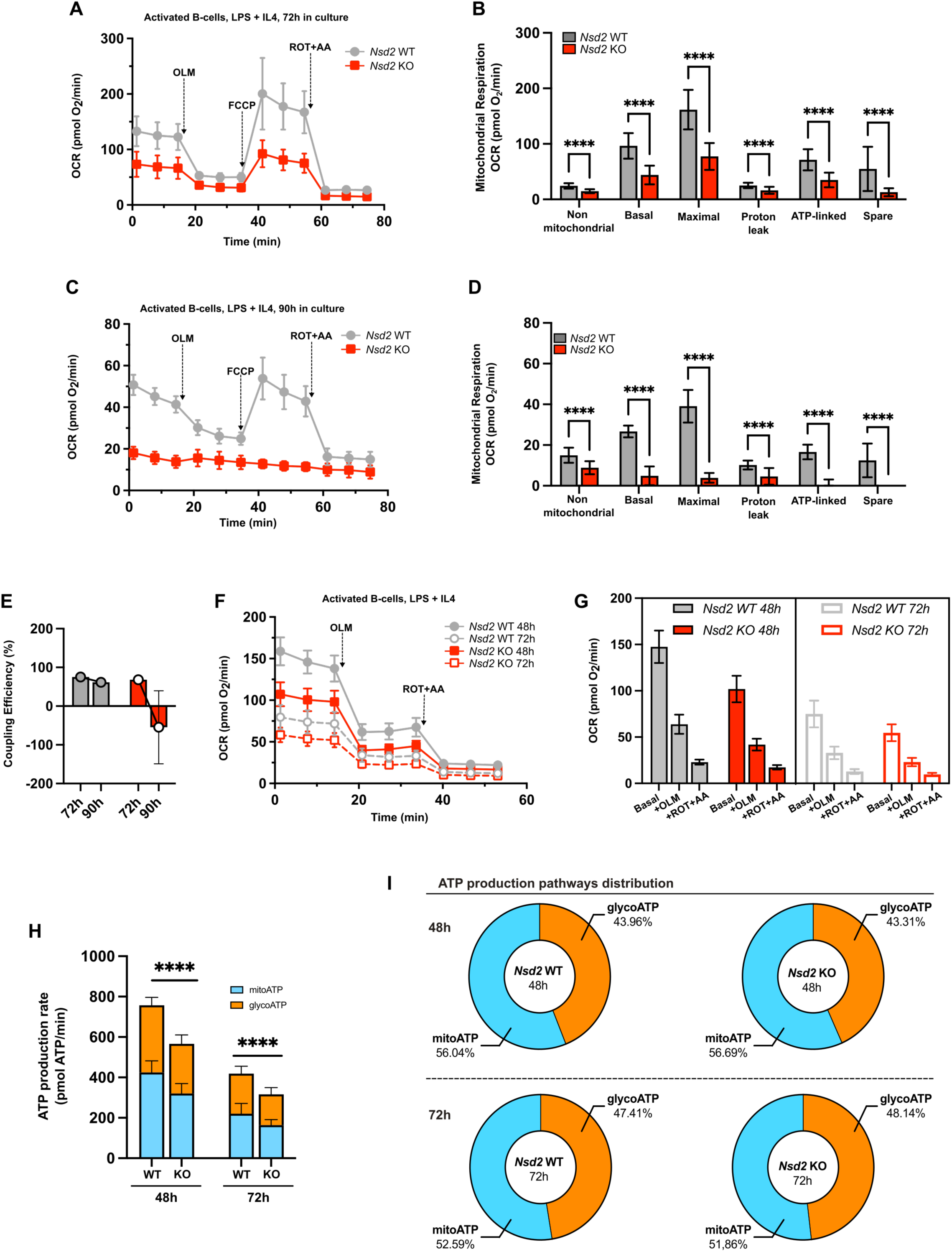
Bioenergetic capacity is decreased in activated *Nsd2^-/-^* B cells. **A-E)** Analysis of mitochondrial respiration in *ex vivo* stimulated B lymphocytes using the *Agilent XF Seahorse Cell Mito Stress* Kit. **A)** Representative graph presenting the data from WT (grey line) and *Nsd2^-/-^* (red line) B lymphocytes cultured for 72 hours in the presence of IL4 and LPS. OLM: oligomycin; FCCP: Carbonyl cyanide-p-trifluoromethoxyphenylhydrazone; ROT+AA: rotenone+antimycin-A. **B)** Main descriptive parameters of mitochondrial respiration in B lymphocyte cultures, calculated from the data shown in panel A. Biological replicates were analyzed in 3 independent experiments with a total of 3 WT and 3 KO mice, with >10 technical replicates per measurement. **C)** Seahorse as in panel A, for cells stimulated 90h in culture. **D)** as in panel B, referred to the data from cells after 90h in culture, from panel C; quantification from 1 experiment with a total of 1 WT and1 KO mice, with 16 technical replicates per measurement. **E)** Coupling efficiency (percentage of basal oxygen consumption dedicated to ATP production in the B cell cultures incubated for the indicated times). **F-I)** Analysis of ATP production in *ex vivo* stimulated B cells using the *Seahorse XF Real-Time ATP Rate Assay* Kit. Biological replicates were analyzed in 2 independent experiments with a total of 3 WT and 3 KO mice, with >30 technical replicates per measurement. **F)** Kinetic profile of Oxygen Consumption Rate measurements in B cells of the indicated genotypes stimulated during the specified times. **G)** Grouped bar graph showing the average OCR values measured at the time points shown in panel F for each condition [Basal respiration (Basal), after addition of oligomycin (+OLM), and after addition of the rotenone + antimycin-A mix (+R/AA)]. **H)** Bar graphs showing ATP production levels derived from glycolysis (orange) or from mitochondrial respiration (blue) for WT and *Nsd2^−/−^* activated B cells, at the indicated time points. **I)** Pie charts representing the relative contribution of glycolysis (orange) and mitochondrial respiration (blue) to total ATP production at 48h and 72h, for WT and *Nsd2^−/−^* activated B cells. Statistical significance was determined using the independent Student’s t-test. *(p<0.05), **(p<0.005), ***(p<0.0005).

Next, we decided to use the Seahorse XF Real-Time ATP kit to determine if the alterations previously detected in the oxygen consumption rate correlated with changes in ATP production in activated *Nsd2^-/-^* B cells. This approach allows distinguishing between the ATP produced via oxidative phosphorylation during mitochondrial respiration (Mitochondrial ATP) and the ATP produced via glycolysis (Glycolytic ATP). Figures 5F-G shows the measurement of OCR in WT and *Nsd2^-/-^* cultures at either 48 or 72 hours after stimulation. Once more, the levels are significantly reduced in knockout versus WT cells at both time points, and in the different conditions after the addition of the inhibitors. Regarding ATP generation, *Nsd2^-/-^* cells show significantly lower levels of ATP production when compared to WT, of both mitochondrial and glycolytic origin, at 48 and 72 hours (Figure 5H), indicating that the loss of Nsd2 function leads to a lower ATP production capacity in B lymphocytes, therefore resulting in a reduction in the energetic potential of the cells. However, this reduction in ATP production affects equally mitochondrial and glycolytic generation, since both are reduced in similar proportions (Figure 5I), with mitochondrial oxidative phosphorylation being the slightly predominant source in both genotypes at either 48h or 72h (Figure 5I).

Given that mitochondrial respiration is contingent on the capacity of processing different types of available substrates, we wanted to study if the aforementioned impairment in the respiration of *Nsd2^-/-^* cells is caused by a lack of capacity for processing some of the main types of substrates. To this aim we used the Agilent XF Substrate Oxidation Stress Test kit, which combines the use of the modulators of cellular respiration with the addition of specific inhibitors of the oxidation pathways of the main primary substrates used by the cell: etomoxir to inhibit the oxidation of long-chain fatty acids; BPTES to inhibit the oxidation of amino acids (glutamine), and UK5099 to inhibit the oxidation of glucose/pyruvate. The results show that basal respiration is not affected by the addition of inhibitors neither in WT nor in *Nsd2^-/-^ ex vivo* activated B cells (Figure 6A). However, all compounds had an effect on the maximum respiratory capacity, to different degrees; the maximum effect was achieved by inhibiting the oxidation of glucose/pyruvate with UK5099, indicating that both cultures are highly dependent on glucose oxidation when cellular energy demand is very high (Figure 6A). In order to more accurately assess the differing impact of these inhibitors on WT versus *Nsd2^-/-^* B cells, we decided to use a normalized version of OCR values, by dividing the values in the presence of inhibitor by the values in its absence (ratio of inhibition = OCR value in the presence of the inhibitor / OCR in its absence, see Figure 6B-F); therefore, a lower ratio indicates a greater impact on the cell’s oxygen consumption due to the inhibition of a specific metabolite utilization. The results show that inhibition of the use of aminoacids doesn’t have a differential impact in *Nsd2^-/-^* versus WT cells (Figure 6D-E) whereas inhibiting the use of glucose (Figure 6F-G) and, especially, of fatty acids (Figure 6B-C), causes a significantly stronger decrease in the oxygen consumption of *Nsd2^-/-^* cells relative to WT ones. These results reveal that while both WT and *Nsd2^-/-^* B cells rely mainly on glucose oxidation under high energy demand, *Nsd2^-/-^* cells show a heightened sensitivity to inhibition of fatty acid and glucose oxidation, suggesting a reduced metabolic flexibility in the absence of Nsd2, pointing to a compromised capacity in *Nsd2^-/-^* cells to efficiently switch between energy substrates to sustain mitochondrial respiration, particularly under stress conditions.

**Figure 6.**
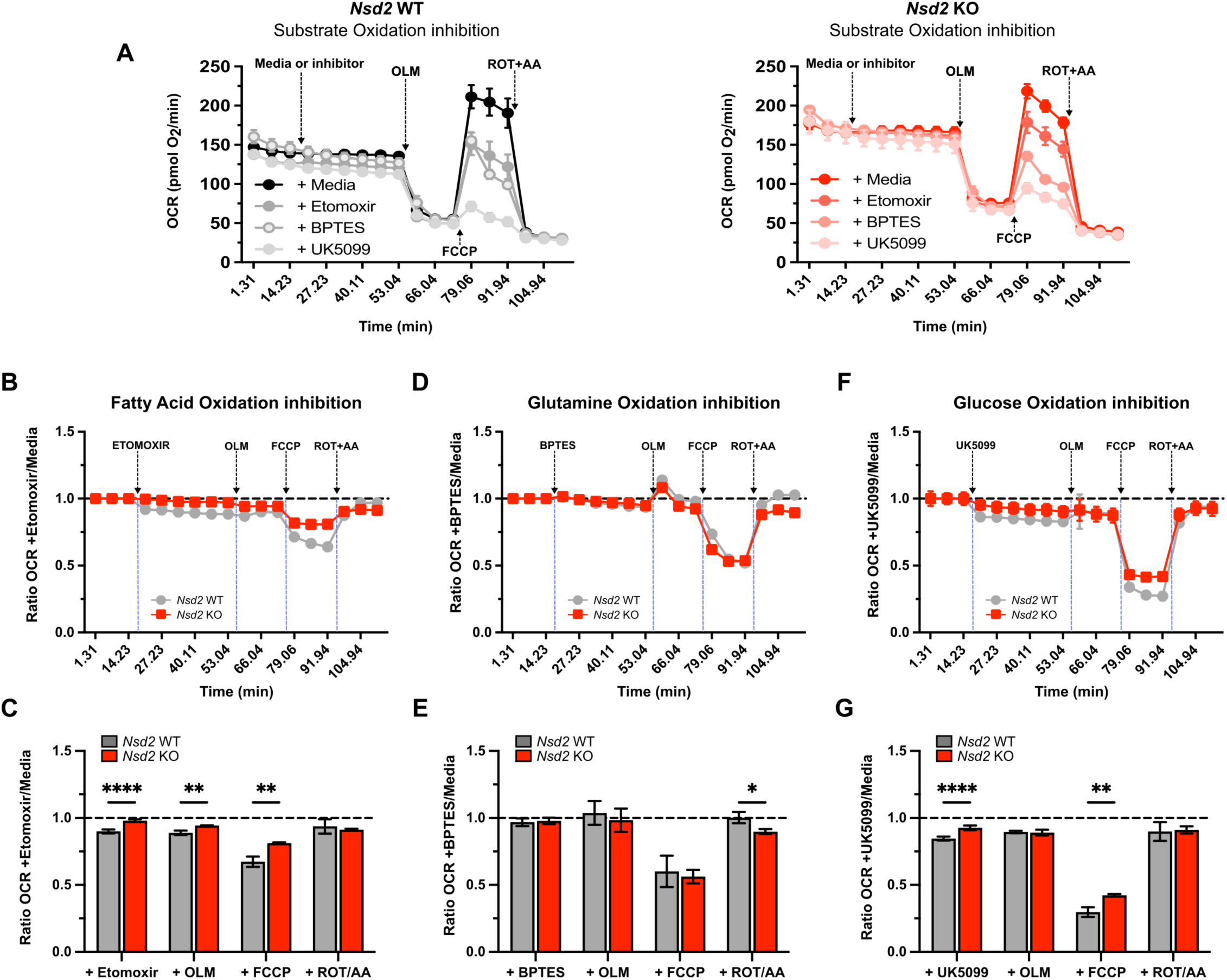
Mitochondrial substrate oxidation profiling reveals reduced metabolic flexibility in *Nsd2^-/-^*B cells. Analysis of mitochondrial respiration in *ex vivo* stimulated B lymphocytes using the Agilent XF Substrate Oxidation Stress Test Kit in the presence of selective inhibitors of the oxidation of the major energy substrates: fatty acids (etomoxir), glutamine (BPTES), and glucose/pyruvate (UK5099). **A)** Kinetic profiles of Oxygen Consumption Rate in wild-type (left, black-grey) and *Nsd2^−/−^* (right, red-pink) stimulated splenic B cells cultured in the presence of the indicated inhibitors. **B-G)** Quantification of the effects of substrate inhibition, shown as the ratio of OCR in the presence of each inhibitor relative to its respective untreated control. Lower ratios reflect greater inhibition of respiration and thus greater dependency on the inhibited substrate. **B-C)** Inhibition of fatty acid oxidation in the presence of etomoxir. **D-E)** Inhibition of amino acid oxidation in the presence of BPTES. **F–G)** Inhibition of glucose/pyruvate oxidation in the presence of UK5099. A total of 400,000 B cells were seeded per well in 180 μl of assay medium. Biological replicates were obtained from a single experiment at 72h: WT (n=1), *Nsd2^−/−^* (n=1), with 15–20 technical replicates per condition. Statistical significance was assessed using an unpaired Student’s t test. *(p<0.05), **(p<0.005), ***(p<0.0005), ****(p<0.0001).

### Nsd2 Loss affects Mitochondrial Function and Structure in Myoblasts and Myotubes

All the aforementioned evidence points to a key role for Nsd2 in regulating mitochondrial function, at least in activated B cells. Since neuromuscular complications (seizures and low muscular tone) are one of the defining core pathological signs of Wolf-Hirschhorn Syndrome, we wanted to ascertain if the loss of *Nsd2* could, just by itself, trigger the mitochondrial alterations that might contribute to this muscular phenotype in WHS patients. To this aim, we first decided to characterize mitochondrial function in *ex vivo* myoblast cultures derived from WT, *Nsd2^+/-^* and *Nsd2^-/-^* mice (Figure 7). One of the advantages of myoblast cultures is that, since they are progenitors, they can be induced to differentiate into myotubes with just a change in the culture medium (Hindi et al., 2017; Yoshioka et al., 2020). Interestingly, Nsd2 was strictly required for myoblast proliferation in culture, in such a way that *Nsd2^-/-^* myoblasts could not be propagated once purified and showed a high percentage of Zombie NIR^+^ cells as an indicator of cell damage (Supplementary Figure 4); in previous research, we have shown that this is also the case for *Nsd2^-/-^* mouse embryo fibroblasts (MEFs), that can be isolated and plated but cannot proliferate in culture beyond the first passage (Campos-Sanchez et al., 2017).

**Figure 7.**
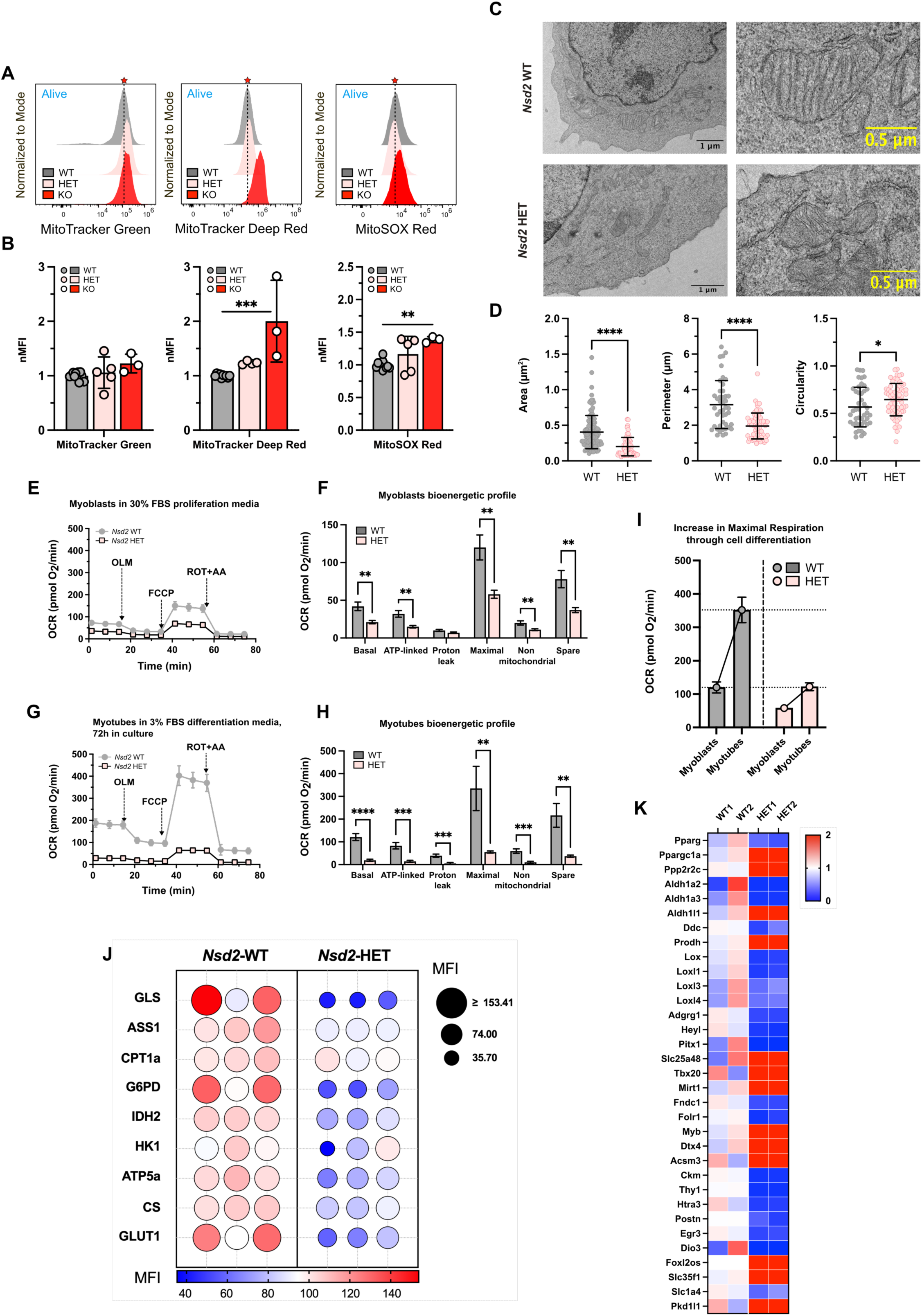
Loss of *Nsd2* alters mitochondrial structure, function, and metabolic gene expression during muscle cell differentiation. **A)** Flow cytometry analysis of mitochondrial mass and function in primary myoblasts from WT, *Nsd2^+/-^*, and *Nsd2^-/-^* mice. Cells were stained with MitoTracker Green (MTG) for mitochondrial mass, MitoTracker Deep Red (MTDR) and TMRM for membrane potential, and MitoSOX Red for mitochondrial ROS. Representative results are shown as normalized mean fluorescence intensity (MFI). Red star and dashed line represent maximum of WT histogram (grey) **B)** Bar graph representing the Mean Fluorescence Intensity of each probe, normalized to the wild-type (WT) control from the corresponding experiment, referred to the data in A. Biological replicates were analyzed from 3 independent experiments: WT (n=3), HET (n=3), and KO (n=5), with 2–3 technical replicates per measurement. Statistical significance was assessed using an unpaired Student’s t test. *(p<0.05), **(p<0.005). **C)** Representative transmission electron microscopy (TEM) images of mitochondria in WT (left) and *Nsd2^+/-^* (right) myoblasts. **D)** Morphometric analysis of mitochondrial area, perimeter, and circularity from TEM images. **E)** Seahorse XF Cell Mito Stress Test showing oxygen consumption rate (OCR) profiles of WT and *Nsd2^+/-^* myoblasts. **F)** Respiratory parameters of WT and *Nsd2^+/-^* myoblasts determined from the data in E. **G)** Seahorse XF Cell Mito Stress Test showing oxygen consumption rate (OCR) profiles of WT and *Nsd2^+/-^* myotubes after 72 of induction of differentiation. **H)** Respiratory parameters of WT and *Nsd2^+/-^* myotubes determined from the data in G. **I)** Increase in OCR following differentiation from myoblasts into myotubes, showing impaired upregulation of mitochondrial function in the transition from *Nsd2^+/-^* myoblasts to myotubes relative to the conversion from WT myoblasts. **J)** Met-Flow profile of three independent biological replicates of WT or *Nsd2^+/-^* myoblasts obtained by the flow cytometry analysis of the levels of the indicated metabolic enzymes. **K)** RNA-seq analysis comparing WT and *Nsd2^+/-^* myoblasts reveals deregulation of genes essential for metabolic rewiring during differentiation. Shown are selected deregulated genes related to mitochondrial metabolism and muscle differentiation.

Contrary to KO cells, heterozygous *Nsd2^+/-^* myoblasts can be grown and propagated in culture, although they were severely reduced in their growth rate in comparison to the WT ones, and they were impaired in their capacity to differentiate into myotubes (data not shown); therefore, we used *Nsd2^+/-^* myoblasts to further characterize the role of Nsd2 in mitochondrial function during muscle development. Using fluorescent probe indicators of mitochondrial function as described above, Figure 7A shows that, while the increase in mitochondrial mass (as determined with MitoTracker Green) is small, both membrane potential (as determined with MTDR) and mitochondrial ROS production (as measured with MitoSOX Red) were significantly increased in *Nsd2^+/-^* and *Nsd2^-/-^* myoblasts when compared with WT cells (Figure 7B), in accordance with the results observed in activated B cells.

To determine whether, similar to B cells, these changes in mitochondrial characteristics measured with the use of fluorescent probes in *Nsd2^+/-^* myoblasts, also correlate with changes in the organelle’s ultrastructure, we inspected these cells using electron microscopy. A representative example of mitochondria from both genotypes is shown in Figure 7C, where it can be appreciated that *Nsd2^+/-^* mitochondria tend to present a smaller area and to have less defined mitochondrial cristae. The graphs in Figure 7D present the results of the measurement of the main parameters defining mitochondrial structure, and they indicate that, similar to activated B cells, mitochondria in *Nsd2^+/-^* myoblasts are significantly smaller than those in WT myoblasts, both in area and perimeter, with altered shape.

Next, we performed extracellular flux analyses to evaluate the mitochondrial respiratory capacity of WT and *Nsd2^+/-^* myoblast cultures; the SeaHorse^TM^ results shown in Figure 7E-F show a clear reduction in the respiratory capacity in *Nsd2^+/-^* myoblasts, manifested in all the different parameters determined, and closely mirroring the previous observations in activated B lymphocytes. It is worth noting that, in the case of B cells, *Nsd2^-/-^* cells, even though they presented a strong mitochondrial phenotype, were viable. However, in the case of myoblasts, this strong phenotype is already present in heterozygous cells, and knockout cells are not viable; this stablishes a closer parallelism with WHS patients, which are hemizygous for *NSD2*. To check whether these functional deficiencies in the mitochondria of *Nsd2^+/-^* myoblasts can also be observed in more advanced stages of muscle development, we induced the differentiation of myoblasts into myotubes, and we determined the mitochondrial respiratory capacity of the differentiated cells. The data in Figure 7G-H show that, also in this more mature cell type, the heterozygous loss of *Nsd2* causes a strong reduction of the oxygen consumption rates, reflected in the significant differences in the respiratory parameters. Indeed, while both genotypes increased oxygen consumption significantly due to the metabolic shift induced by differentiation, the increase in OCR of *Nsd2^+/-^* myotubes was significantly lower compared to the WT ones (Figure 7I), further widening the gap in oxygen consumption between the two genotypes under mitochondrial stress.

Met-Flow (Ahl et al., 2020) is a flow cytometry-based method that allows examining the metabolic state of the cells by measuring the expression levels of key proteins across the metabolic network. We have used this method (Figure 7J) to compare the levels of expression of several critical proteins between WT and *Nsd2^+/-^* myoblasts: Glutaminase (GLS, responsible of catalyzing the hydrolysis of glutamine), Argininosuccinate Synthase 1 (ASS1, involved in aminoacid synthesis), Carnitine palmitoyltransferase I (CPT1a, essential for the mitochondrial uptake and beta-oxidation of long-chain fatty acids), Glucose-6-phosphate dehydrogenase (G6PD, a key enzyme of the pentose phosphate pathway), NADP(+)-dependent mitochondrial isocitrate dehydrogenase (IDH2, essential in the TCA cycle), Hexokinase I (HK1, the first step in most glucose metabolism pathways), ATP Synthase F1 Subunit Alpha (ATP5a, a subunit of the mitochondrial ATP synthase), Citrate Synthase (CS, essential for TCA cycle), and Glucose Transporter Type 1 (GLUT1, a major glucose transporter). The data in Figure 7J show that all these essential proteins are significantly downregulated in different biological replicates of *Nsd2^+/-^* myoblasts, highlighting the metabolic impairment occurring in muscle progenitors when the dose of this epigenetic regulator is decreased.

These results indicate that Nsd2 plays a critical role in maintaining mitochondrial integrity in muscle precursor cells, not only by supporting respiratory capacity but also by preserving proper mitochondrial architecture. The fact that such profound mitochondrial dysfunction is already evident in heterozygous *Nsd2^+/-^* cells underscores the sensitivity of muscle cells to Nsd2 dosage, and reinforces the hypothesis that mitochondrial impairment caused by *NSD2* hemizygosis may contribute to the neuromuscular manifestations observed in Wolf-Hirschhorn Syndrome.

### Loss of Nsd2 impairs mitochondrial remodeling in muscle differentiation/ commitment

In view of the fact that the reduction of the dosage of *Nsd2* in myoblasts largely recapitulates the mitochondrial and respiratory phenotypes seen in *Nsd2^-/-^* stimulated B cells, we wanted to study how these alterations might be related to changes in gene expression. To this aim, we performed RNAseq-based differential gene expression analysis comparing WT vs. *Nsd2^+/-^* myoblasts (since *Nsd2^-/-^* myoblasts do not grow in culture) (Figure 7K). The data show a general deregulation (either down or up) of key genes involved in the correct rewiring of metabolism necessary to supply the energetic needs of differentiating muscle cells (Figure 7K), like Creatine Kinase (*Ckm*), Peroxisome Proliferator-Activated Receptor Gamma (*Pparg*), Peroxisome Proliferator-Activated Receptor Gamma Coactivator 1 Alpha (*Pgc-1α*), aldehyde dehydrogenases *Aldh1a2*, *Aldh1a3* and *Aldh1l1*, Dopa decarboxylase (*Ddc*), Proline Dehydrogenase (*Prodh*), Mitochondrial Ribosomal Translation Factor 1 (*Mirt1*), Folate Receptor 1 (*Folr1*), or Acyl-CoA Synthetase Medium Chain Family Member 3 (*Acsm3*). This deregulation of metabolism-related genes happens in the context of a general impairment of *Nsd2^+/-^* myoblasts to activate the gene program required to develop into muscle cells, with a downregulation of genes like *Myb*, *Pitx1*, *Heyl*, *Tbx20*, *Postn*, Lysyl Oxidases *Lox*, *Loxl1*, *Loxl3*, *Loxl4*, and others (Figure 7K). These findings support a critical role for Nsd2 in activating the metabolic program required for successful differentiation into myotubes.

### Nsd2 Loss affects Mitochondrial Structure in Adult Skeletal and Cardiac Muscle

In a mixed C57Bl/6J x 129sv background, *Nsd2^-/-^* mice are viable and can grow to adulthood, although they are significantly smaller than their WT littermates (Nimura et al., 2009). The results obtained in cultured myoblasts showed that Nsd2 is necessary for correct muscle differentiation from progenitors. This fact, together with the loss of muscular tone affecting the majority of WHS patients, prompted us to study if the loss of Nsd2 might also affect mitochondrial structure in adult muscular tissue in *Nsd2^-/-^* mice. To this aim, we analyzed by TEM samples of both skeletal and cardiac muscle of adult *Nsd2^-/-^* animals. A representative example of the images captured for skeletal muscle is depicted in Figure 8A; the main two types of muscular mitochondria that can be observed in muscle fibers are clearly distinguishable: subsarcolemmal mitochondria (SSM, adjacent to the membrane), and interfibrillar mitochondria (IFM, located between the different myofibrils). In *Nsd2^-/-^* skeletal muscle cells, SSM are slightly smaller than in WT cells, but there are no relevant changes in their perimeter or shape (Figure 8B). However, the morphology of interfibrillar mitochondria was more severely affected by the loss of *Nsd2*, with significantly smaller size and perimeter when compared with mitochondria from WT cells (Figure 8C).

**Figure 8.**
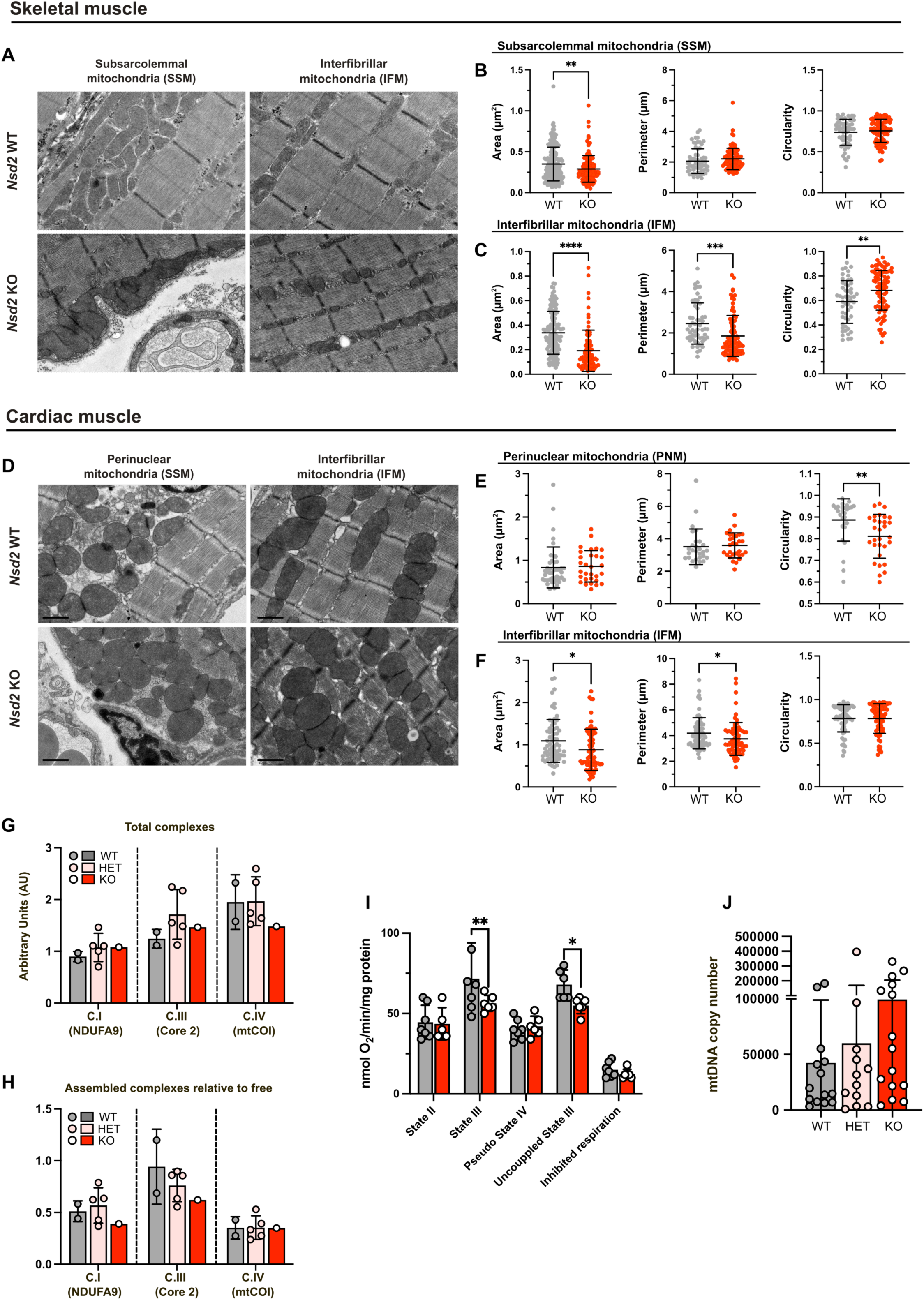
*Nsd2* deficiency alters mitochondrial ultrastructure and impairs mitochondrial function in adult skeletal and cardiac muscle. **A)** Transmission electron microscopy (TEM) images of skeletal muscle from adult WT and *Nsd2^-/-^* mice. Two distinct mitochondrial populations are highlighted: subsarcolemmal mitochondria (SSM), located beneath the sarcolemma, and interfibrillar mitochondria (IFM), situated between myofibrils. **B)** Morphometric analysis of SSM mitochondria in WT and *Nsd2^-/-^* skeletal muscle based on TEM images. Parameters measured include mitochondrial area, perimeter, and circularity. **C)** Morphometric quantification of IFM mitochondria from WT and *Nsd2^-/-^* skeletal muscle. Significant reductions in mitochondrial area and perimeter are observed in *Nsd2^-/-^* samples. **D)** Representative TEM images of cardiac muscle from WT and *Nsd2^-/-^* mice. Perinuclear mitochondria (PNM) and interfibrillar mitochondria (IFM) are identified. **E)** Morphometric quantification of PNM mitochondria from WT and *Nsd2^-/-^* cardiomyocytes. **F)** Morphometric quantification of IFM mitochondria from WT and *Nsd2^-/-^* cardiomyocytes. Significant reductions in mitochondrial area and perimeter are observed in *Nsd2^-/-^* samples. **G-H)** Blue Native PAGE (BN-PAGE) analysis of respiratory chain complex assembly in mitochondria isolated from skeletal muscle of WT, *Nsd2^+/-^*, and *Nsd2^-/-^* mice. Immunoblotting was performed using antibodies against representative subunits of Complexes I-IV and measured by densitometry (see Supplementary Figure 5); G) Quantification shows reduced total levels of Complex III and IV, and H) decreased assembly of Complex I and III into supercomplexes in *Nsd2^-/-^* samples. **I)** Mitochondrial respiration assays using a Clark-type oxygen electrode on isolated mitochondria from WT and *Nsd2^-/-^* soleus muscle. **J)** Quantification of mitochondrial DNA (mtDNA) copies in total muscle tissue from WT, *Nsd2^+/-^*, and *Nsd2^-/-^* mice. Biological replicates were analyzed from a total of 3 WT, 3 HET and 3 KO mice, with 3 technical replicates per measurement. *(p<0.05), **(p<0.005).

In the case of cardiomyocytes (Figure 8D), mitochondria are classified into perinuclear mitochondria (PNM, adjacent to the cell nucleus and with the main function of supplying energy for gene transcription), and interfibrillar mitochondria (IFM, found between myofibrils and supplying energy for myocardial contraction). Again in this case, electron microscopy revealed small differences in PNM (Figure 8E), while IFM in *Nsd2^-/-^* cardiomyocytes presented significant changes in their morphology (Figure 8F), being smaller in both size and perimeter when compared with interfibrillar mitochondria in WT cardiomyocytes. These results suggest that the lack of Nsd2 function translates into alterations in mitochondrial morphology in both skeletal and cardiac muscle tissue, with a particularly relevant reduction in mitochondrial size, being the extent of these alterations dependent on the intracellular localization of the mitochondria.

To check if these morphological defects correlated with alterations in mitochondrial function, mitochondria were freshly isolated from skeletal muscle from WT, *Nsd2^+/-^* and *Nsd2^-/-^* animals (Fernandez-Silva et al., 2007), and their respiratory chain complexes and supercomplexes where analyzed using Blue Native Polyacrylamide Gel Electrophoresis (BN-PAGE) (Wittig et al., 2006) and immunoblotted with specific antibodies against the different subunits of the respiratory complexes (Figure 8G-H and Supplementary Figure 5). The results show that, in *Nsd2^-/-^* muscle, the amount of total Complex III and Complex IV is reduced in comparison to WT (Figure 8G) and that, particularly for Complexes I and III, the amount of assembled complexes relative to free ones is notably reduced in the skeletal muscle from *Nsd2^-/-^* animals (Figure 8H). Since this decreased assembly might lead to reduced mitochondrial efficiency, to evaluate this aspect we performed mitochondrial respiration analyses using a Clark-type oxygen electrode (Sanchez-Gonzalez and Formentini, 2021) in isolated mitochondria from mouse skeletal muscle (Figure 8I); the results revealed a significant reduction in ADP-stimulated (State III) and uncoupled (FCCP-induced) respiration in skeletal muscle mitochondria from *Nsd2^-/-^* mice compared to WT controls. These findings indicate a specific impairment in oxidative phosphorylation capacity and in maximal electron transport system function in the absence of *Nsd2^-/-^*, further supporting a critical role for this gene in achieving mitochondrial bioenergetic efficiency in adult muscle.

Finally, to further characterize the effect of *Nsd2* deficiency in mitochondrial function in adult muscle, we quantified the amount of mitochondrial DNA (Cao et al., 2022) in extracts of muscle from mice of the three genotypes (Figure 8J), and we could observe clear differences showing an increase in the number of copies of mtDNA as the dosage of *Nsd2* is reduced. This increase in mtDNA may represent a compensatory response to impaired mitochondrial function, since it has been described that, under conditions of mitochondrial dysfunction, cells often activate pathways that promote mitochondrial biogenesis in an attempt to restore bioenergetic homeostasis (Bernal-Tirapo et al., 2023; Filograna et al., 2019; Filograna et al., 2021).

All these data support a role for Nsd2 in the correct maintenance of mitochondrial structure and bioenergetic function in adult muscle tissue, and further link Nsd2 deficiency to the muscular phenotypes observed in WHS patients.

## DISCUSSION

Epigenetic regulators are in charge of re-defining cellular identity during differentiation, a process that requires the integration of multiple cellular tasks, among others the coordination between transcriptional regulation and metabolic adaptation. In several tissues and developmental stages, metabolic reprogramming is a strict requirement to fuel the energy-demanding processes. Two examples of such developmental crossroads are terminal B cell development and early muscle differentiation, both cell types whose function is compromised in Wolf-Hirschhorn Syndrome patients. As an H3K36me2 methyltransferase, NSD2 has been recognized for its role in cell identity maintenance (Hoetker et al., 2023), and emerging evidence suggests that H3K36 methylation controls lineage plasticity at developmental transitions (Hoetker et al., 2023; Pashos et al., 2025; Sussman et al., 2025), with loss of NSD2 leading to extensive gene expression changes that impair correct differentiation. In different types of tumors, the ectopic overexpression of NSD2 has been shown to cause different types of metabolic alterations affecting distinct routes, although disentangling these effects from other signaling cascades is not easy in the context of tumoral cells (Chong et al., 2023; Sobh et al., 2024; Wang et al., 2016). In any case, its role in controlling the reprogramming of metabolism during normal, physiological, developmental processes is still unclear.

B cell activation requires profound metabolic adaptations to support clonal expansion and antibody production (Chen et al., 2021a). Recent research has shown that regulation of metabolic pathways in activated immune cells plays a critical role in fate decision and function (Ganeshan and Chawla, 2014; Jang et al., 2015; Jung et al., 2019; Pearce and Pearce, 2013; Waters et al., 2018; Weisel et al., 2020). Naïve B cells are quiescent, while GC B cells (as activated B cells) are highly proliferative and metabolically active (Haniuda et al., 2020; Iborra-Pernichi et al., 2024; Mendoza et al., 2018; Weisel et al., 2020). Activated B cells at the GC undergo metabolic rewiring in response to stimulation, which seems coupled with an increase in both mitochondria-dependent and -independent metabolism (Haniuda et al., 2020; Iborra-Pernichi et al., 2024; Mendoza et al., 2018; Weisel et al., 2020). We had previously shown that the loss of *Nsd2* interferes with the induction of the transcriptional activation and signaling components that control the process of GC formation (Campos-Sanchez et al., 2017), while also impairing correct DNA replication, leading to inefficient repair of DNA damage and a reduction of cellular fitness. The results presented here show that, beyond the control over the transcriptional activators in charge of the specification of cellular identity, Nsd2 also plays a key role regulating a plethora of nuclear genes involved, at many levels, in the activation of the mitochondrial and metabolic reprogramming necessary to satisfy the energetic demands of these processes. Nsd2 loss perturbs oxidative phosphorylation (OXPHOS), mitochondrial function, and cellular metabolism. RNA-seq analysis of *Nsd2^-/-^* activated B cells revealed deregulation of mitochondrial respiratory complexes, metabolic regulators, and chromatin-associated factors that influence metabolism. Supporting this, Seahorse metabolic analyses revealed that Nsd2-deficient B cells exhibit impaired oxygen consumption rates (OCR) and ATP production, indicative of metabolic inflexibility.

GC B cells, contrary to naïve B cells, minimally utilize aerobic glycolysis and instead use fatty acids to conduct oxidative phosphorylation (Weisel et al., 2020). To dissect the metabolic consequences of *Nsd2* loss, we assessed substrate utilization in B cells. Our findings reveal that Nsd2-deficient cells exhibit increased sensitivity to fatty acid and glucose oxidation inhibition in comparison to WT cells, indicating an impaired ability to adapt to metabolic demands. Together with the deregulation of key metabolic enzymes, these results suggest that Nsd2 is necessary for balancing energy sources during differentiation. Additionally, our *in vivo* and *ex vivo* studies confirm that *Nsd2* loss leads to mitochondrial membrane hyperpolarization and excessive ROS accumulation, which correlates with increased cellular stress and reduced cellular fitness. It has recently been shown that the enzyme with the opposite function to Nsd2 (the H3K36me2 demethylase Kdm2a) needs to be downregulated in activated B cells for the process of affinity maturation to take place correctly (Nakagawa et al., 2024); indeed, inhibition of Kdm2a enhances OXPHOS by optimizing the expression of key nuclear mitochondrial genes, while preventing excessive production of reactive oxygen species (ROS) (Nakagawa et al., 2024). Now, our data support the fact that a precise control of H3K36me2 levels during activation of B cells, mediated by an Nsd2/Kdm2a balance, is essential for the regulated expression of nuclear mitochondrial genes and adequate mitochondrial remodeling, which are, in turn, necessary for the survival and proliferation of the cells. A similar antagonism has been described for these epigenetic modifiers in the regulation of epithelial-mesenchymal transitions in cancer (Yuan et al., 2020); loss of KDM2A enhances fatty acid oxidation and metabolic reprogramming (Chen et al., 2021b), whereas *NSD2* deficiency shifts cells toward metabolic inefficiency. These findings highlight an H3K36 methylation-dependent regulatory axis that fine-tunes mitochondrial metabolism and cellular fitness.

Similar to B cells, muscle progenitors also undergo a metabolic shift during differentiation, transitioning from a proliferative glycolytic state to an oxidative mitochondrial metabolism (Bobori et al., 2023; Pereira et al., 2011; Sin et al., 2016). In adipocyte development, it has previously been shown that the expression of a mutated form of histone H3 than cannot be methylated in position 36 interferes with the correct activation of the genes required for adipocyte differentiation, and also affects the expression of myogenic genes in the C2C12 cell line (Zhuang et al., 2018). Now, our results show that the metabolic reprogramming mediated by Nsd2 is essential for myogenic differentiation, with Nsd2 haploinsufficiency leading to defective myoblast commitment and ineffective differentiation into myotubes.

Consistent with its role in B cells, a decrease in *Nsd2* levels in myoblasts resulted in impaired mitochondrial respiration, increased ROS levels, and decreased ATP production. Furthermore, differentiating *Nsd2^+/-^* myotubes failed to effectively upregulate mitochondrial activity, reinforcing the idea that Nsd2 is required for metabolic adaptation during differentiation. TEM analysis further confirmed that mitochondria in *Nsd2*-deficient myoblasts were smaller and structurally compromised, mirroring the ultrastructural defects observed in B cells. Given that H3K36 methylation is crucial for lineage stability (Hoetker et al., 2023; Pashos et al., 2025), our findings suggest that Nsd2 safeguards muscle differentiation by coordinating chromatin modifications with mitochondrial remodeling. In agreement with this, the skeletal muscle of adult *Nsd2^-/-^* mice presents alterations in mitochondrial distribution as determined by electron microscopy, together with an abnormal assembly of the respiratory complexes in the inner mitochondrial membrane, and an impairment in oxidative phosphorylation capacity. Also, *Nsd2^-/-^* muscle fibers present an altered ratio of mtDNA to genomic DNA, which might be caused by a compensatory increase in mitochondrial content in an attempt of the cells to counterbalance a reduced oxidative phosphorylation capacity caused by a compromised mitochondrial function (Bernal-Tirapo et al., 2023; Filograna et al., 2019; Filograna et al., 2021).

The interplay between epigenetics and metabolism is increasingly recognized as a fundamental mechanism governing cell fate. H3K36 methylation modulates gene expression, DNA methylation, and enhancer accessibility, providing a stable framework for lineage commitment (Brumbaugh et al., 2019; Hoetker et al., 2023). Our data establish Nsd2 as a key integrator of epigenetics and metabolism, a central regulator of chromatin-modulated metabolic reprogramming that is crucial for immune cell activation and muscle differentiation, and whose loss of function results in widespread metabolic gene deregulation and mitochondrial dysfunction. By linking epigenetic modifications with mitochondrial function, Nsd2 ensures the proper execution of metabolic transitions required for cell fate determination. In the context of Wolf-Hirschhorn Syndrome, our results reaffirm the key role of NSD2 hemizygosis in the patient’s symptoms, and suggest that its partial loss alone may suffice to cause both the neuromuscular and the immune defects observed in the patients. Furthermore, our results support the hypothesis that mitochondrial dysfunction contributes to the multi-organ pathology observed in affected individuals.

Given that metabolic interventions are increasingly being explored as therapeutic strategies in diseases characterized by mitochondrial dysfunction (El-Hattab et al., 2017; Guo et al., 2013; Wallace, 2018), our results raise the possibility that targeting metabolic pathways could help alleviate some of the developmental and functional deficits observed in Wolf-Hirschhorn Syndrome patients. Ultimately, given that metabolic rewiring is a common feature of lineage plasticity and cancer progression (Yuan et al., 2020), a deeper understanding of how NSD2 orchestrates metabolic rewiring during differentiation may not only offer insight into disease mechanisms but also open avenues for therapeutic modulation of metabolism in both tumoral and developmental epigenetically-driven disorders.

## EXPERIMENTAL PROCEDURES

Detailed Materials and Methods are listed at the end of the *Supplementary Information* document.

## ACKNOWLEDGEMENTS

The authors thank all the members of their laboratories for discussion. Research at C. Cobaleda’s laboratory was partially supported by Grant PID2021-122787OB-I00 funded by MICIU/AEI/ 10.13039/501100011033 and by “ERDF/EU”, by the Fundación Científica de la Asociación Española contra el Cáncer (PRYCO211305SANC), by the Fundacion Unoentrecienmil (CUNINA2 project), and by a Research Contract with the “Fundación Síndrome de Wolf-Hirschhorn o 4p-”. Sara Cogliati was a recipient of a ‘Ramón y Cajal fellowship 23013-2017’ founded by MCIN/AEI/10.13039/501100011033 and ‘El FSE invierte en tu futuro’. Research in Sara Cogliati’s lab is supported by the Grant PID2020-114054RA-I00 1001100482 founded by MCIN/AEI/10.13039/501100011033. Research in Nuria Martínez-Martín lab is partially supported by HR22-00447 from La Caixä, and PID2021-126298OB-100 financed by the Ministerio de Ciencia, Innovación y Universidades. Institutional grants from the “Fundación Ramón Areces” and “Banco de Santander” to the CBMSO are also acknowledged. The CBM is a Severo Ochoa Center of Excellence (CEX2021-001154-S), funded by MCIN/AEI/10.13039/501100011033. J. Martínez-Cano was partially supported by a predoctoral fellowship FPI-UAM 2019. B.S. Estrada is supported by PRE2022-102696 from the Ministerio de Ciencia, Innovación y Universidades. We also thank Drs. Cristina Sanchez-González and Laura Formentini for their help with measurements with the Clark’s electrode. We thank members of the CBM Electron Microscopy, Flow Cytometry and Animal House facilities for their support and advice.

## AUTHOR CONTRIBUTIONS

Initial conception of the project was designed by C.C. and J.M.-C.. J.M.-C conducted most experiments and data acquisition, developed most of the methodology and was in charge of maintenance, management and supervision of the mouse colony. BN-PAGE and mitochondrial DNA content experiments were performed by A.S. and supervised by S.C. and C.C. MetFlow experiments were performed by J.M-C. and B.S.E. and supervised by N.M.-M and C.C. J.M.-C. and C.C. were responsible for analysis and interpretation of the data. J.M.-C. assembled the figures. Funding acquisition, supervision and project administration, C.C.. Editing of draft, J.M.-C., S.C., N.M.-M. and C.C. All authors gave feedback on the paper.

## DECLARATION OF INTERESTS

The authors declare no competing interests.

## Notes

### Competing Interest Statement

The authors have declared no competing interest.

