## Supplementary_Figures_and_Methods for "NSD2 links developmental plasticity to mitochondrial function in muscle and lymphocyte differentiation"

[1] *Immune system development and function* Unit, Centro de Biología Molecular Severo Ochoa (Consejo Superior de Investigaciones Científicas - Universidad Autónoma de Madrid), Madrid, Spain.
[2] Program of *Physiological and Pathological Processes*, Centro de Biología Molecular Severo Ochoa (Consejo Superior de Investigaciones Científicas - Universidad Autónoma de Madrid), Madrid, Spain.
[3] Program of *Tissue and Organ Homeostasis*, Centro de Biología Molecular Severo Ochoa (Consejo Superior de Investigaciones Científicas - Universidad Autónoma de Madrid), Madrid, Spain.
[4] *Intestinal Morphogenesis and Homeostasis* Group, Area 3-Cancer, Instituto Ramón y Cajal de Investigación Sanitaria (IRYCIS), Madrid, Spain

**KEYWORDS:** Wolf-Hirschhorn Syndrome; Nsd2; Whsc1; Metabolic Reprogramming; Epigenetic Plasticity; Mitochondrial Function; Immunodeficiency; Neuromuscular defects.

**RUNNING TITLE:** NSD2 COUPLES EPIGENETICS TO MITOCHONDRIAL FUNCTION

**#Correspondence should be addressed to:**
César Cobaleda,
*Immune system development and function Unit, Centro de Biología Molecular Severo Ochoa (CSIC- UAM), Laboratory 325, Calle Nicolás Cabrera 1, 28049, Madrid, Spain.*
https://www.cbm.uam.es/ccobaleda
vs.2025 04 26

#### SUPPLEMENTARY FIGURES

**Supplementary Figure 1. Increased percentage of activated B lymphocytes with dysfunctional mitochondria in the absence of Nsd2, *ex vivo* and *in vivo*. A)** Representative FACS plots showing the analysis of mitochondrial membrane potential (as measured with MitoTracker Deep Red) vs cell division (as measured by dilution of CellTrace) in *ex vivo* activated WT and *Nsd2*$^{-/-}$ B lymphocytes. Cells with dysfunctional mitochondria appear as MitoTracker Depp Red$^{LOW}$ and CellTrace$^{HIGH}$ (undivided). B cells were incubated for 72 hours in proliferation medium supplemented with IL-4 and LPS. Biological replicates were analyzed in three independent experiments with 3 WT mice and 5 *Nsd2*$^{-/-}$ mice. Each measurement included 2–3 technical replicates. **B)** Representative FACS plots showing the analysis of mitochondrial membrane potential (as measured with MitoTracker Deep Red) vs. mitochondrial mass (as measured by MitoTracker Green) in splenocytes from WT and *Nsd2*$^{-/-}$ mice, 7 days after immunization with Sheep Red Blood Cells. Cells with dysfunctional mitochondria appear as MitoTracker Depp Red$^{LOW}$ and MitoTracker Green$^{HIGH}$. **C)** Bar Graph representing the percentage of Cells with dysfunctional mitochondria for the experiments shown in panels A and B. Statistical significance was determined using an unpaired Student’s t-test. *$p < 0.05$, **$p < 0.005$, ***$p < 0.0005$, ****$p < 0.0001$.

**Supplementary Figure 1**

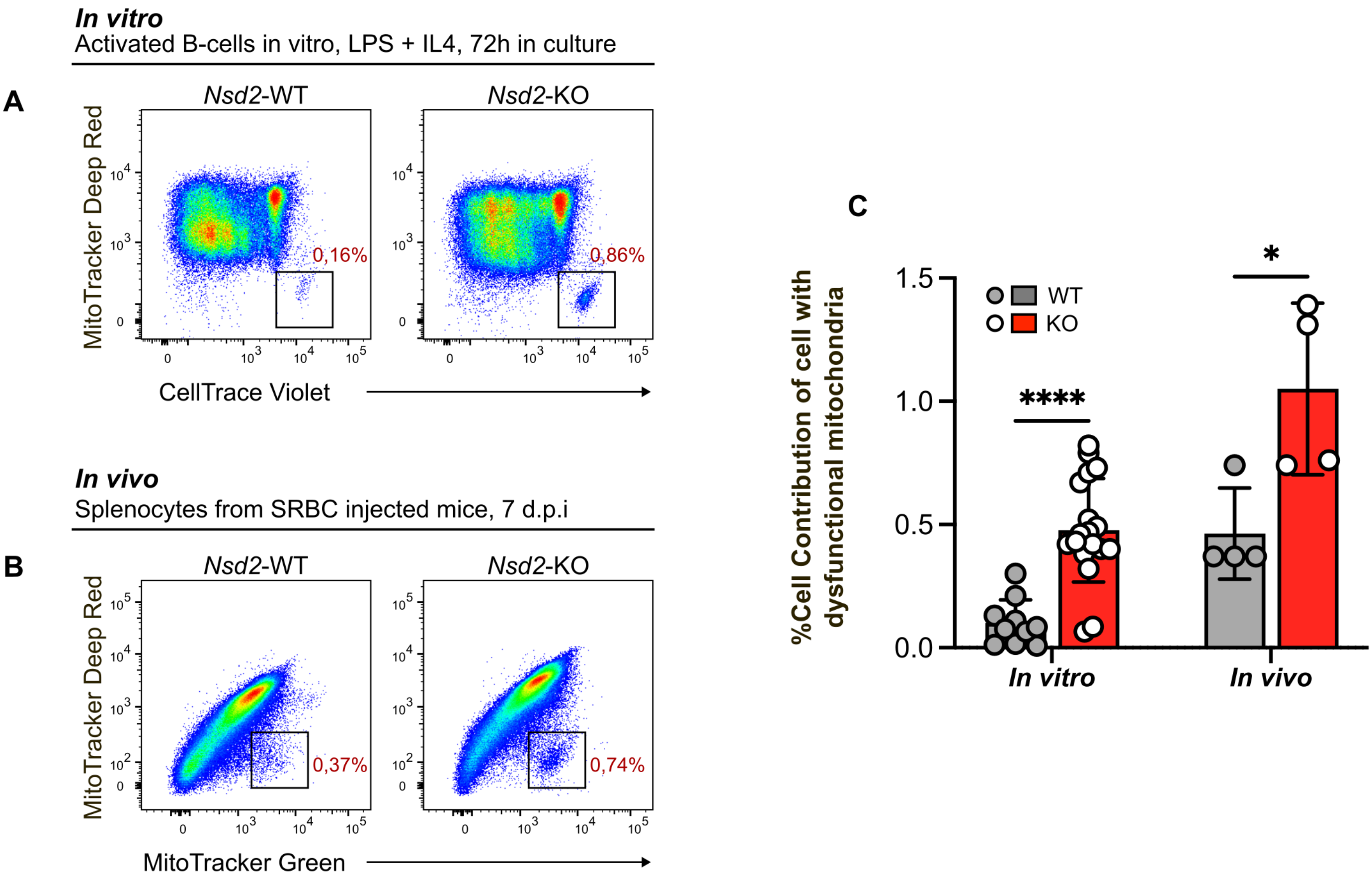

**Supplementary Figure 2. Variations in mitochondrial mass across generations in *ex vivo* activated *Nsd2*-deficient B lymphocytes as determined by flow cytometric analysis in the presence of CellTrace Violet and MitoTracker Green**. **A)** Representative example of flow cytometry analysis from an *ex vivo* B cell activation assay after 72 hours of culture in the presence of IL-4 and LPS. FACS plots show the number of cell divisions (as indicated by dilution of CellTrace Violet fluorescence peaks). Comparison of *Nsd2*$^{+/-}$ and *Nsd2*$^{-/-}$ profiles versus WT one, normalized to mode. G0-G5: generations (cell divisions) in culture, as deconvoluted with FlowJo software. Accumulation of undivided cells (G0) is evident in KO cells. **B)** Percentage of cell contribution relative to WT for each generation in culture, for *Nsd2* heterozygous and knockout cells. **C-H)** Bar graphs of normalized MitoTracker Green MFI values for each generation, referenced to the corresponding WT control (grey bar) for the three cell subpopulations indicated [total culture (left, panels C,F), IgG1$^{-}$ cells (center, panels D,G), and IgG1$^{+}$ cells (right, panels E,H)] for *Nsd2*$^{+/-}$ cells (panels C-E, pink columns) and *Nsd2*$^{-/-}$ cells (panels F-H, red columns). B cells were incubated for 72 hours in proliferation medium containing IL-4 and LPS. Biological replicates analyzed from three independent experiments: WT (n=3 mice), *Nsd2*$^{+/-}$ (n=3), *Nsd2*$^{-/-}$ (n=5). Each measurement included 2–3 technical replicates.

#### Supplementary Figure 2

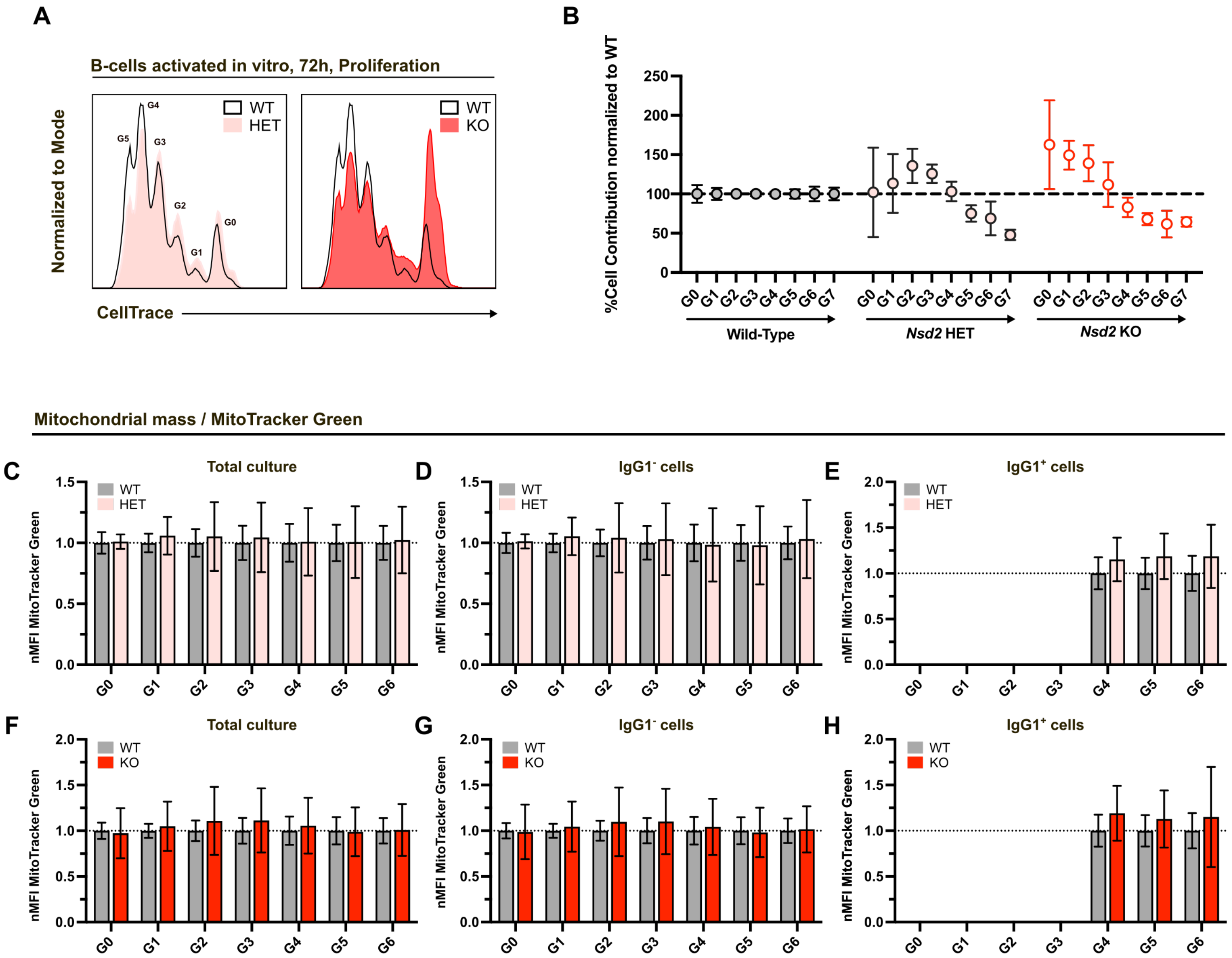

**Supplementary Figure 3. Time-dependent decline in the bioenergetic capacity of activated *Nsd2*$^{-/-}$ B cells**. Time-course analysis of mitochondrial respiration of *ex vivo* stimulated B lymphocytes using the *Agilent XF Seahorse Cell Mito Stress* Kit. **A)** General scheme of the time-course experiment: MACS-purified splenic B cells from mice of the indicated genotypes (WT –grey color– or *Nsd2*$^{-/-}$ –red color–) were incubated in proliferation medium containing IL-4 and LPS for the indicated times. At each time point, a SeaHorse analysis was performed and the different respiratory parameters determined. **B-G)** Bar graph representations of the evolution of the indicated respiratory parameters with time in culture, for WT and KO activated B cells. Biological replicates analyzed from three independent experiments: WT (n=3 mice), *Nsd2*$^{-/-}$ (n=5). Each measurement included 2–3 technical replicates. Statistical significance was assessed using an unpaired Student's t test. *(p<0.05), **(p<0.005), ***(p<0.0005), ****(p<0.0001).

#### Supplementary Figure 3

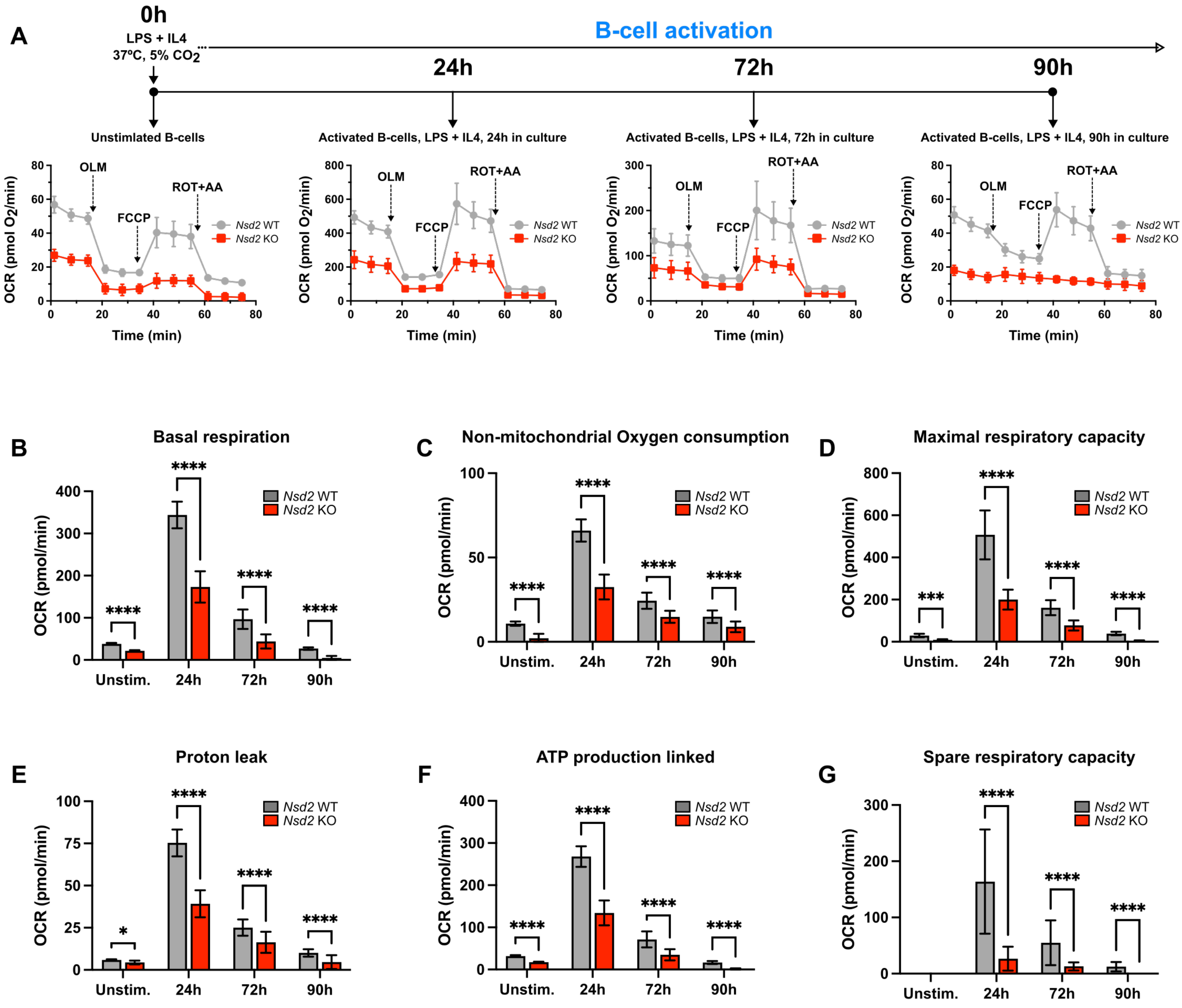

**Supplementary Figure 4. Skeletal myoblasts derived from *Nsd2*$^{-/-}$ mice cannot proliferate in culture.** FACS analysis of freshly stablished myoblast cultures for the indicated genotypes [Solid dark grey, WT; solid pink, *Nsd2*$^{+/-}$ (HET); solid red, *Nsd2*$^{-/-}$ (KO)] **A)** *Upper row*: expression of the myoblast marker Integrin-7α in the stablished culture. *Lower row:* identification of dead or damaged myoblasts using Zombie NIR staining as an indicator of cell damage. **B)** Quantification of living cells and cells in the Integrin-7α$^{+}$ myoblast compartment for each genotype. Statistical significance was assessed using an unpaired Student's t test. *($p<0.05$), **($p<0.005$), ***($p<0.0005$), ****($p<0.0001$).

### Supplementary Figure 4

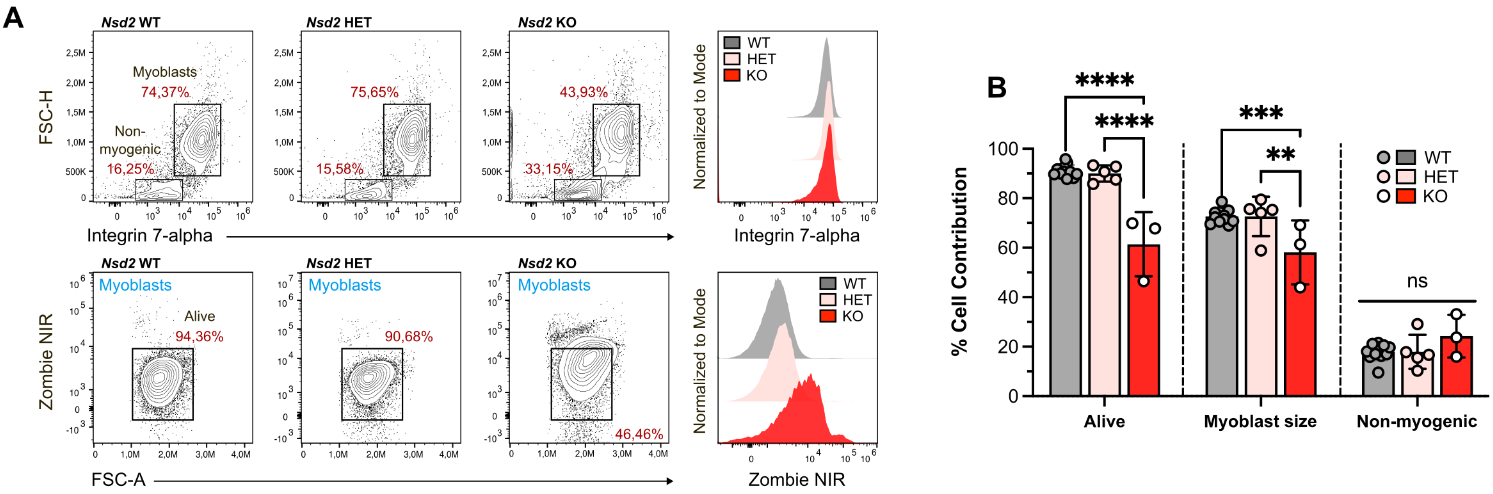

**Supplementary Figure 5. Effects of Nsd2 deficiency on the assembly of mitochondrial supercomplexes in skeletal muscle. A)** Blue Native PAGE of mitochondria isolated from skeletal muscle of mice of the indicated genotypes. **B-L)** Immunodetection of the indicated subunits of the different mitochondrial Complexes: B-C) Ndufa9 (CI), D-E) Core2 (CIII), F) Ndufa9+Core2 merge, G-H) COI (CIV), I) Ndufa9+Core2+COI merge, J-K) VDAC, L) Ndufa9+Core2+COI+VDAC merge.

#### Supplementary Figure 5

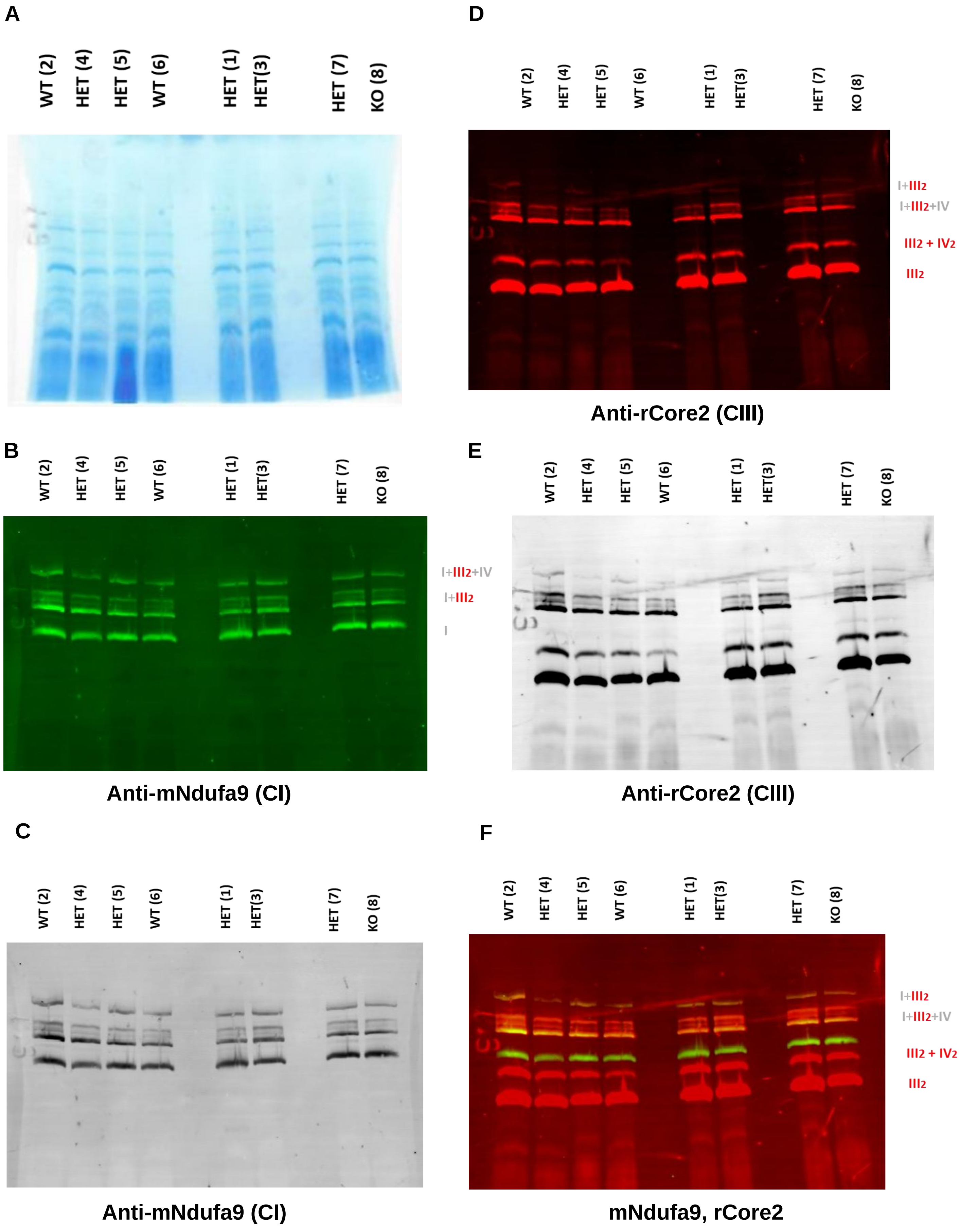

#### Supplementary Figure 5

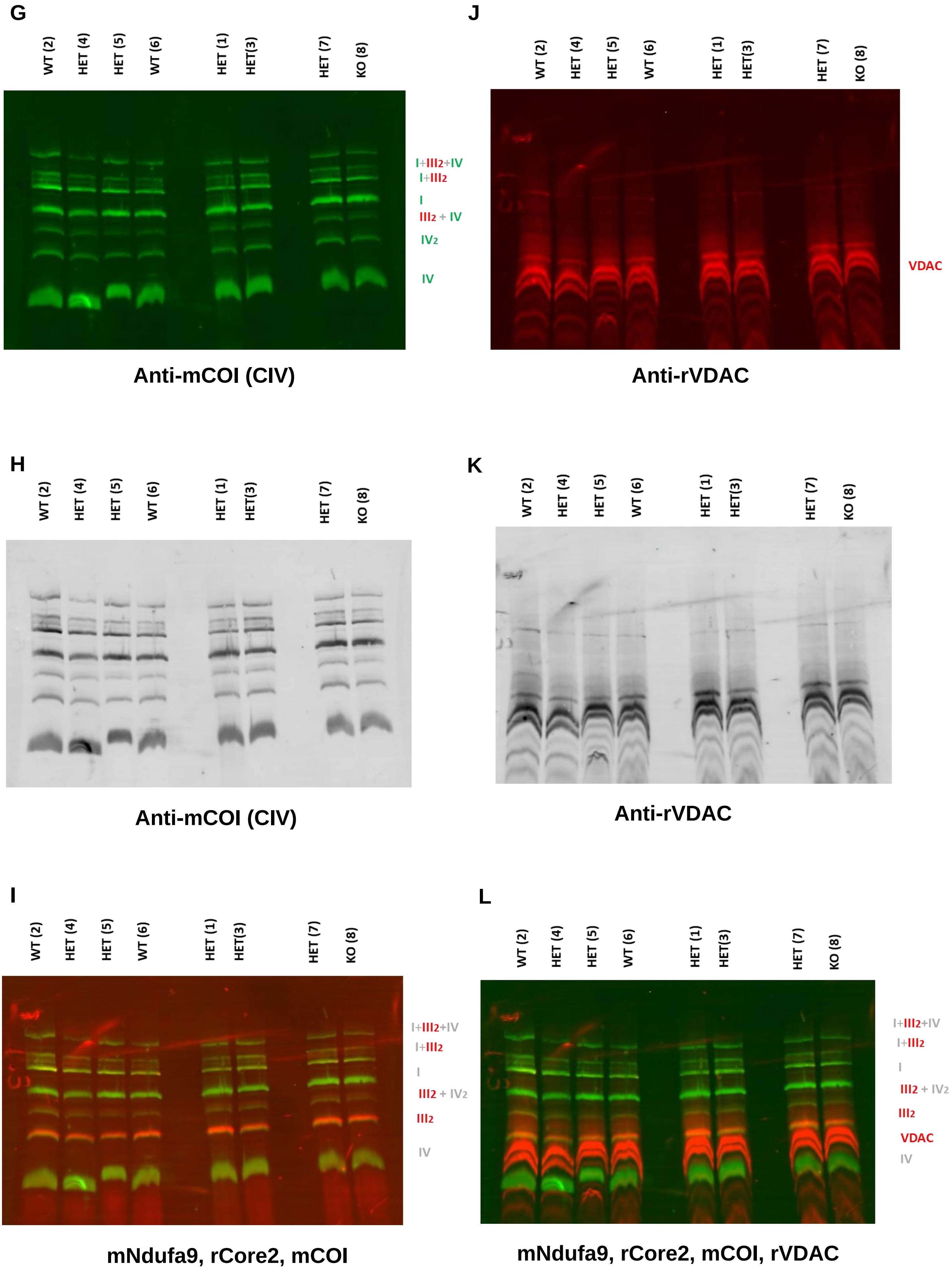

#### EXPERIMENTAL PROCEDURES

**Mice.** *Nsd2$^{-/-}$* mice were maintained in a hybrid C57BL/6 x 129/SvJ background (since the KO is lethal in a pure C56BL/6 background) and, when required, genotyped as described (Nimura et al., 2009). All mice were housed in a special pathogen-free (SPF) barrier facility. All experimental procedures conducted on mice were approved by the CBM's Institutional and Spanish Regional Committees on Ethics of animal research. Mice with the indicated genotypes were included in the study without any further preselection or formal randomization and comprised balanced numbers from both genders; we used age-matched mice. Investigators were not blinded to genotype group allocations.

**FACS Analysis and Sorting Purification.** Mice were humanely sacrificed and single cell suspensions were prepared from organs in FACS buffer (1X PBS with 2% heat-inactivated fetal bovine serum [HI-FBS, Thermo Fisher Scientific]). Contaminating red blood cells were lysed using ACK Lysing Buffer (0.15 M NH4Cl, 10 mM KHCO3, 0.1 mM EDTA; pH = 7.2-7.4). The remaining cells were washed in FACS buffer. After staining, all cells were washed once in FACS buffer and resuspended in the same buffer containing 2 mg/ml propidium iodide (PI) to allow dead cells to be excluded. In addition, Zombie NIR Fixable Viability Dye (BioLegend) was used as live/dead cell discriminator at a 1:5000 dilution. The samples and the data were acquired in a FACSCantoII Flow Cytometer or Cytek Aurora full spectrum cytometer and analyzed using FlowJo software (TreeStar). Preincubation of cells with CD16/CD32 (2.4G2) Fc-block solution (BD Biosciences) was used to avoid nonspecific antibody binding. The different conjugated antibodies used for surface staining of the cells in this study are indicated in Table 1.

**B-cell isolation, labelling and activation assays**. Mouse primary B cells were purified from the spleens of the indicated strains of mice by immunomagnetic depletion with anti-CD43 beads (Miltenyi Biotec) and were cultured in 50 µM 2-β-Mercaptoethanol (Invitrogen), 10 mM Hepes (Invitrogen), 1 mM Glutamine, antibiotic mix (penicillin and streptomycin, Thermo Fisher Scientific), Non-Essential Aminoacids and 10% HI-FBS RPMI medium, supplemented with 25 µg/ml LPS (Sigma-Aldrich) and 20 ng/ml IL4 (PeproTech). CellTrace$^{TM}$ Violet Cell Proliferation Kit (Invitrogen) was used as proliferation tracer following manufacturer's instructions. Mitochondrial probes are listed in Table 2.

**Sheep Red Blood Cell (SRBC) immunizations.** Mice were immunized with 200 µl/mouse of SRBCs (Oxoid) resuspended in 1X PBS at a concentration of $10^9$ cells/ml, by intraperitoneal injection. SRBCs were washed on 1X PBS at least 3 times before counting. Spleens of immunized mice were extracted at the indicated time points after SRBC injection.

**Isolation, Culture, and Differentiation of Primary Myoblasts from Adult Mouse Skeletal Muscle.** Primary myoblasts were isolated from the anterior tibial, quadriceps, and gastrocnemius muscles of adult mice as previously described (Hindi et al., 2017; Shahini et al., 2018; Yoshioka et al., 2020), with some modifications. Muscles were dissected post-sacrifice, cleaned of fat, minced, and digested for 1 h at 37°C in DMEM (Thermo Fisher Scientific) containing 400 U/ml collagenase II (Worthington) and antibiotic mix. Digestion was neutralized using DMEM + 5% HI-FBS, filtered first through 70 µm and immediately after through 40 µm cell strainers, and cells were pelleted by centrifugation and resuspended in proliferation medium (DMEM + 20% HI-FBS + 10% horse serum [Thermo Fisher Scientific] + 1% chicken embryo extract [MP Biomedicals] + 10 ng/ml bFGF [PeproTech], 50 µg/ml gentamicin [Merck] + 2 µg/ml ciprofloxacin [Sigma]). Cells were seeded in 6-well culture plates coated with Matrigel (Corning) diluted to 10% (v/v) in DMEM. The coating was performed under sterile conditions in a biosafety cabinet. Each well was completely covered with the diluted Matrigel solution and incubated at room temperature for 30 seconds. After this brief incubation, the excess Matrigel was aspirated, and the plates were left uncovered for approximately 20 minutes under sterile airflow to allow the surface to dry. Plates were then maintained at room temperature inside the biosafety cabinet until cell seeding. Proliferating medium was replaced every 48 h. Upon reaching confluence, myoblasts were either purified via pre-plating—brief incubation in uncoated plates to deplete non-myogenic cells—or passaged to 10 cm Matrigel-coated plates. For passaging, cells were trypsinized (0.25% Trypsin-EDTA [Merck], 3 min at 37°C), centrifuged, and resuspended in fresh proliferation medium. Cells were either replated or frozen in proliferation medium supplemented with 10% DMSO (Sigma). To induce differentiation, confluent myoblast cultures (85–95%) were switched to differentiation medium (DMEM + 5% FBS + antibiotic mix) and maintained at 37°C, 5% $CO_2$ until myotube formation was evident upon microscope examination.

**Met-flow cytometry staining**

According to (Ahl et al., 2020), nine metabolic enzymes were selected and optimized for flow cytometry analysis of primary myoblast cultures. Myoblasts were obtained from previously described primary cultures and maintained in proliferation medium until the time of the experiment. A single-cell suspension was generated by trypsinization, as previously described. Two antibodies, ASS1-AF647 (1:2500) and IDH2-PE (1:2500), were purchased pre-conjugated. The remaining antibodies were conjugated in-house using purified forms and appropriate fluorochromes: G6PD-PECy7 (1:2500), HK1-AF647 (1:100), ATP5A-AF594 (1:500), CPT1A-AF488 (1:100), CS-AF488 (1:1000), GLS-AF700 (1:1000), and GLUT1-AF405 (1:100). Surface staining was performed with integrin α7-APC (1:100), and Zombie NIR was used as a viability marker. Extracellular staining was carried out at 4 °C for 20 minutes in Brilliant Stain Buffer (BD, 563794) diluted in PBS with 2% HI-FBS. After staining, cells were washed, then fixed and permeabilized using the Foxp3/Transcription Factor Staining Buffer Set (eBioscience), following the manufacturer's instructions. Cells were subsequently washed in the provided wash buffer and incubated in wash buffer supplemented with 20% HI-FBS for 30 minutes at room temperature to block nonspecific binding. Intracellular staining was performed in permeabilization buffer for 2 hours at room temperature. Cells were then washed once with permeabilization buffer and once with PBS containing 2% HI-FBS. Samples were acquired on a Cytek Aurora 5-laser spectral cytometer, and data were analyzed using FlowJo software.

**Extracellular flux assay (SEAHORSE)**

Metabolic flux analyses were performed using the Seahorse XF96 Extracellular Flux Analyzer (Agilent Technologies) on two distinct cell types: in vitro activated, purified B lymphocytes and primary myoblasts, obtained as previously explained. Cells were seeded in Seahorse XF96 cell culture microplates pre-coated with either 10% Matrigel (for myoblasts) or 22.4 µg/mL Cell-Tak (Corning, 354240) (for B cells). Assays were conducted using the Mito Stress Test, ATP Rate Assay, and Substrate Oxidation Stress Test kits (Agilent Technologies), following the manufacturer's protocols. For the assays, cells were resuspended in Seahorse XF RPMI medium (for B cells) or Seahorse XF DMEM medium (for myoblasts), both supplemented with 10 mM glucose, 2 mM pyruvate, and 20 mM glutamine. Oxygen consumption rate (OCR) data were normalized per cell using a Cytation 1/5 plate reader (BioTek) and Gen5 software v3.11 (BioTek). For B lymphocytes, normalization was performed based on high-contrast bright field image-based cell counting. For myoblasts, nuclei were stained with 10 µM Hoechst 33342 (Thermo Fisher Scientific) prior to image acquisition and quantification.

**RNA Extraction, Sequencing and Bioinformatic Analyses.** Myoblast RNA was extracted following commercial Trizol® Reagent (Life Tecnologies™) recommendations. Its chemical quality was measure by Nanodrop 1000 Spectrophotometer (Thermo Fisher Scientific) and its integrity was confirmed using the Agilent 2100 Bioanalyzer. RNA samples were quality-controlled and processed by Novogene for transcriptomic analysis. mRNA was enriched using poly-T oligo-attached magnetic beads and fragmented prior to cDNA synthesis. Library preparation followed a non-strand-specific protocol and included end repair, A-tailing, adapter ligation, PCR amplification, and purification. Libraries were quantified using Qubit and qPCR, assessed by Bioanalyzer for fragment size distribution, and sequenced on an Illumina platform. Raw sequencing reads were quality-filtered using fastp, and clean reads were aligned to the mouse reference genome using HISAT2 v2.0.5 (Mortazavi et al., 2008). Transcript assembly was performed with StringTie v1.3.3b (Mortazavi et al., 2008), and gene expression quantification was conducted using featureCounts v1.5.0-p3 (Liao et al., 2014), generating FPKM values for each gene. Differential expression analysis between genotypes was performed with the DESeq2 R package (Love et al., 2014), applying Benjamini–Hochberg correction for multiple testing. Gene Set Enrichment Analysis was performed with GSEA (Broad Institute) (Subramanian et al., 2005), using the data obtained after normalization and quantification. The use of the whole data matrix instead of preranked data allows the obtention of heatmaps for the different genesets employed, as shown in Figure 1.

**Accession numbers**

Accession numbers for RNA-seq data are GEO GSE84878 and GSE88970. Other data are available from the authors upon request.

**Statistical analysis.**

Exact sample sizes and statistical tests used for comparisons are indicated in each figure. Kaplan–Meier survival curves were created and analyzed using Prism (GraphPad Software), which used the log-rank test to calculate significance. Samples were allocated to their experimental groups according to their predetermined type (i.e., mouse genotype) and, therefore, there was no randomization. Sample sizes chosen are indicated in the individual figure legends and were not based on formal power calculations to detect pre-specified effect sizes. Data analysis was not blinded.

#### Transmission Electron Microscopy (TEM)

For ultrastructural analysis, cell and tissue samples were processed for TEM following standard procedures. Cultured cells (e.g., myoblasts, activated B cells) and mouse tissue samples (soleus skeletal muscle and heart) were chemically fixed in 4% paraformaldehyde and 2% glutaraldehyde in 0.1 M PBS (pH 7.4), followed by post-fixation in 1% osmium tetroxide with 0.8% potassium ferricyanide, and contrasted with 0.15% tannic acid and 2% uranyl acetate. After ethanol dehydration, samples were embedded in TAAB 812 epoxy resin (TAAB Laboratories). For tissue samples, propylene oxide was used as a transitional solvent before resin infiltration. Polymerization was carried out at 60°C for 48 hours. Ultrathin sections (60–70 nm) were cut using a Leica Ultracut UCT ultramicrotome and mounted on Formvar-coated copper-palladium grids. Sections were contrasted with 2% aqueous uranyl acetate and Reynolds' lead citrate before imaging. TEM analysis was performed using a JEOL JEM-1010 electron microscope operated at 80 kV. Digital micrographs were acquired using a 4K × 4K TVIPS F416 CMOS camera. Mitochondrial classification and mitochondrial area were segmented manually and analysed using Fiji (http://fiji.sc/Fiji) and ImageJ 1.48v software

#### Mitochondria Isolation

Mitochondria were isolated from gastrocnemius muscle following the previously described protocol (Fernandez-Silva et al., 2007). Briefly, skeletal muscle tissue was minced with a scalpel blade and homogenized in a buffer containing 0.32 M sucrose, 1 mM EDTA, and 10 mM Tris-HCl (pH 7.4), using a Potter-Elvehjem homogenizer at 600 rpm. Homogenates were subjected to a low-speed centrifugation at 600 ×g for 10 minutes at 4 °C to remove debris. The supernatant was collected and centrifuged twice at 10,000 ×g for 10 minutes at 4 °C to pellet mitochondria. The resulting mitochondrial pellet was resuspended in the same sucrose-based buffer. Protein concentration was determined using the Bradford assay (Bio-Rad).

#### Mitochondrial Membrane Solubilization

To analyze respiratory chain complexes and supercomplexes, isolated mitochondria were resuspended at 10 mg/mL in solubilization buffer containing 1.5 M aminocaproic acid and 50 mM Bis-Tris/HCl (pH 8.0). Digitonin was added to a final concentration of 4%, and samples were incubated on ice for 5 minutes to solubilize mitochondrial membranes. Lysates were centrifuged at 16,000 ×g for 20 minutes at 4 °C, and the supernatant was collected. Prior to electrophoresis, samples were mixed with Serva Blue G dye (Serva Electrophoresis, 5% stock solution).

#### Blue Native Gel Electrophoresis and Immunoblotting

Respiratory chain complexes were separated under non-denaturing conditions by Blue Native Gel Electrophoresis (BNGE) using a 3–12% gradient gel as previously described (Wittig et al., 2006). Gels were prepared using a peristaltic pump and cast into a Mini-PROTEAN system (Bio-Rad). After electrophoresis, proteins were transferred onto Hybond-P polyvinylidene fluoride (PVDF) membranes (GE Healthcare). Immunoblotting was performed using primary antibodies specific for subunits of the individual respiratory complexes (see Table 2). Secondary antibodies conjugated to Alexa Fluor 680 or Alexa Fluor 800 (Invitrogen) were used for detection. Fluorescent signals were acquired using the ODYSSEY Infrared Imaging System (LI-COR).

**Mitochondrial DNA content**

Mitochondrial DNA content was measured as described previously (Cao et al., 2022) with some modifications. DNA extractions from muscle were performed following instructions in DNeasy kit (QIAGEN), and using 100µl of elution buffer in the last step. Nuclear and mitochondrial DNA were amplified by quantitative PCR with 10 ng of total DNA as template, using specific primers for mitochondrial gene mtCOI (mtCO1 Forward: CTGAGCGGGAATAGTGGGTA; mtCO1 Reverse: TGGGGCTCCGATTATTAGTG; nSDHA Forward: TACTACAGCCCCAAGTCT), and nuclear gene SDHA (nSDHA Forward: TACTACAGCCCCAAGTCT; nSDHA Reverse: T TGGACCCATCTTCTATGC), and the GoTaq(R) qPCR Master mix (Promega). Mitochondrial content was normalized to nuclear DNA, and was calculated using the equation: $\Delta C_T$ = (nucDNA $C_T$ – mtDNA $C_T$). Relative mitochondrial DNA copies = $2 \times 2^{\Delta CT}$

**Respiration Assays Using Clark Electrode for Mitochondrial State Analysis**

Oxygen consumption assays were performed using a Clark electrode-based Oxytherm-R respirometer (HansTech) to assess mitochondrial respiration across different states. Assays were conducted according to the protocol described by (Sanchez-Gonzalez and Formentini, 2021). Fresh mitochondria were isolated as described above. For each assay, 300 µg of isolated mitochondria were used. The substrates employed included glutamate-malate (final concentration 10 mM), which facilitates electron entry through Complex I of the electron transport chain (ETC); succinate (final concentration 10 mM), which donates electrons to Complex II; and malate in combination with palmitoyl-carnitine (final concentrations 0.5 mM and 0.5 mM, respectively), which allows for the coupling of oxygen consumption to β-oxidation. The analysis aimed to assess different mitochondrial respiratory states, including state 2 (baseline), state 3 (maximal ADP-stimulated respiration), and state 4 (non-phosphorylating respiration).

**Table 1.** Conjugated antibodies used for surface FACS staining

| Antigen | Clone | Fluorochrome | Provider | Reference | Dilution |
|---|---|---|---|---|---|
| B220 | RA3-6B2 | FITC | BDBioscience | 553088 | 1:500 |
| B220 | RA3-6B2 | APC | BioLegend | 553092 | 1:500 |
| Cd19 | 6D5 | Pacific Blue | BDBioscience | 115523 | 1:500 |
| Cd95 | Jo2 | PE | BDBioscience | 554258 | 1:500 |
| GL7 | GL7 | Biotin | BDBioscience | 13.5902-82 | 1:500 |
| IgG1 | A85-1 | Brilliant Violet 605 | BDBioscience | 563285 | 1:500 |
| Integrin 7a | 3C12 | APC | Miltenyi Biotec | 130-123-833 | 1:100 |
| Streptavidin | - | APC-Cy 7 | BDBioscience | 554063 | 1:1000 |
| CD16/CD32 FcBlock | 2.4G2 | Purified | BDBioscience | 553142 | 1:100 |

**Table 2**. Other reagents

| *Antibodies* | | |
|---|---|---|
| **Reagent or resource** | **Provider** | **Reference** |
| Anti-G6PD Clone EPR 20668 | Abcam | ab210702 |
| Anti-Glut1 Clone EPR 3915 | Abcam | ab210438 |
| Anti-ASS1 Clone EPR 12398 | Abcam | ab170952 |
| Anti-ATP5A Clone EPR13030 | Abcam | ab176569 |

| | | |
|---|---|---|
| Anti-Hexokinase1 Clone EPR10134 | Abcam | ab150423 |
| Anti-CPT1A Clone 8F6AE9 | Abcam | ab128568 |
| Anti-IDH2 Clone EPR7577 | Abcam | ab131263 |
| Anti-GLS Clone EPR28716-72 | Abcam | ab317032 |
| Anti-CS Clone EPR8067 | Abcam | ab129095 |
| Anti-UQCRC2 (Core2). Host: rabbit | Protein Tech | 14742-1-AP |
| Anti-VDAC1. Host: rabbit | Abcam | Ab15895 |
| Anti-NDUFA9. Host: mouse | Abcam | Ab14713 |
| Anti-MTCO1. Host: mouse | Invitrogen | 459600 |
| Anti-RABBIT 680. Goat anti-rabbit IgG | Invitrogen | A21076 |
| Anti-MOUSE 800. Goat anti-mouse IgG | Invitrogen | SA5-10176 |
| *Chemical, peptides and recombinant proteins* | | |
| **Reagent or resource** | **Provider** | **Reference** |
| CD43 (Ly48) MicroBeads, mouse | Miltenyi Biotec | 130-049-801 |
| CellTrace Violet | Thermo Fisher Scientific | C34557 |
| Lipopolysaccharides from E. coli (LPS) | Sigma-Aldrich | L2630 |
| Mouse Interleukin 4 (IL4) Recombinant protein | PeproTech | 214-14-1MG |
| Albumin Fraction V (ph 7.0) | PanReac AppliChem | A1391,0250 |
| Matrigel Matrix | Corning | 356234 |
| Chicken Embryo Extract | MP Biomedicals | 092850145 |
| Human FGF-basic (FGF-2/bFGF) Recombinant protein | PeproTech | 100-18B-1MG |
| *Viability probes for Flow Cytometry* | | |
| **Reagent or resource** | **Provider** | **Reference** |
| Propidium Iodide (PI) | Sigma | P4170 |
| Zombie NIR Fixable Viability Kit | BioLegend | 423105 |
| *Mitochondrial probes for Flow Cytometry* | | |
| **Reagent or resource** | **Provider** | **Reference** |
| MitoTracker Green FM | Thermo Fisher Scientific | M7514 |
| MitoTracker Deep Red | Thermo Fisher Scientific | M22425 |
| Tetramethylrhodamine, methyl ester, perchlorate (TMRM) | Thermo Fisher Scientic | T668 |
| MitoSOX Red | Thermo Fisher Scientific | M36008 |
| *Agilent's Consumables for Seahorse technology* | | |
| **Reagent or resource** | **Provider** | **Reference** |
| Seahorse XF Cell Mito Stress Test Kit | Agilent | 103010-100 |
| Seahorse XF Cell Real-Time ATP Rate Assay kit | Agilent | 103591-100 |
| Seahorse XF Long Chain Fatty Acid Oxidation Stress Test Kit | Agilent | 103672-100 |
| Seahorse XF Glucose/Pyruvate Oxidation Stress Test Kit | Agilent | 103673-100 |

| | | |
|---|---|---|
| XF Glutamine Oxidation Stress Test kit | Agilent | 103674-100 |
| Seahorse XF DMEM Medium, pH 7.4 | Agilent | 103575-100 |
| Seahorse XF RPMI Medium, pH 7.4 | Agilent | 103576-100 |
| Seahorse XF 1.0 M Glucose | Agilent | 103577-100 |
| Seahorse XF 100 mM Pyruvate | Agilent | 103578-100 |
| Seahorse XF 200 mM Glutamine | Agilent | 103579-100 |

#### METHODS' REFERENCES


Ahl, P.J., Hopkins, R.A., Xiang, W.W., Au, B., Kaliaperumal, N., Fairhurst, A.M., and Connolly, J.E. (2020). Met-Flow, a strategy for single-cell metabolic analysis highlights dynamic changes in immune subpopulations. Commun Biol *3*, 305.

Cao, Y., Vergnes, L., Wang, Y.C., Pan, C., Chella Krishnan, K., Moore, T.M., Rosa-Garrido, M., Kimball, T.H., Zhou, Z., Charugundla, S., *et al.* (2022). Sex differences in heart mitochondria regulate diastolic dysfunction. Nat Commun *13*, 3850.

Fernandez-Silva, P., Acin-Perez, R., Fernandez-Vizarra, E., Perez-Martos, A., and Enriquez, J.A. (2007). In vivo and in organello analyses of mitochondrial translation. Methods Cell Biol *80*, 571-588.

Hindi, L., McMillan, J.D., Afroze, D., Hindi, S.M., and Kumar, A. (2017). Isolation, Culturing, and Differentiation of Primary Myoblasts from Skeletal Muscle of Adult Mice. Bio Protoc *7*.

Liao, Y., Smyth, G.K., and Shi, W. (2014). featureCounts: an efficient general purpose program for assigning sequence reads to genomic features. Bioinformatics *30*, 923-930.

Love, M.I., Huber, W., and Anders, S. (2014). Moderated estimation of fold change and dispersion for RNA-seq data with DESeq2. Genome Biol *15*, 550.

Mortazavi, A., Williams, B.A., McCue, K., Schaeffer, L., and Wold, B. (2008). Mapping and quantifying mammalian transcriptomes by RNA-Seq. Nat Methods *5*, 621-628.

Nimura, K., Ura, K., Shiratori, H., Ikawa, M., Okabe, M., Schwartz, R.J., and Kaneda, Y. (2009). A histone H3 lysine 36 trimethyltransferase links Nkx2-5 to Wolf-Hirschhorn syndrome. Nature *460*, 287-291.

Sanchez-Gonzalez, C., and Formentini, L. (2021). An optimized protocol for coupling oxygen consumption rates with beta-oxidation in isolated mitochondria from mouse soleus. STAR Protoc *2*, 100735.

Shahini, A., Vydiam, K., Choudhury, D., Rajabian, N., Nguyen, T., Lei, P., and Andreadis, S.T. (2018). Efficient and high yield isolation of myoblasts from skeletal muscle. Stem Cell Res *30*, 122-129.

Subramanian, A., Tamayo, P., Mootha, V.K., Mukherjee, S., Ebert, B.L., Gillette, M.A., Paulovich, A., Pomeroy, S.L., Golub, T.R., Lander, E.S., *et al.* (2005). Gene set enrichment analysis: a knowledge-based approach for interpreting genome-wide expression profiles. Proc Natl Acad Sci U S A *102*, 15545-15550.

Wittig, I., Braun, H.P., and Schagger, H. (2006). Blue native PAGE. Nat Protoc *1*, 418-428.

Yoshioka, K., Kitajima, Y., Okazaki, N., Chiba, K., Yonekura, A., and Ono, Y. (2020). A Modified Pre-plating Method for High-Yield and High-Purity Muscle Stem Cell Isolation From Human/Mouse Skeletal Muscle Tissues. Front Cell Dev Biol *8*, 793.
